# Asparagine availability controls B cell homeostasis

**DOI:** 10.1101/2023.04.03.535433

**Authors:** Yavuz F. Yazicioglu, Eros Marin, Hana F. Andrew, Karolina Bentkowska, Julia C. Johnstone, Robert Mitchell, Zhi Yi Wong, Kristina Zec, Joannah Fergusson, Mariana Borsa, Iwan G. A. Raza, Moustafa Attar, Mohammad Ali, Barbara Kronsteiner, Izadora L. Furlani, James I. MacRae, Michael J. Devine, Mark Coles, Christopher D. Buckley, Susanna J. Dunachie, Alexander J. Clarke

## Abstract

Germinal centre (GC) B cells proliferate at some of the highest rates of any mammalian cell, yet the metabolic processes which enable this are poorly understood. We performed integrated metabolomic and transcriptomic profiling of GC B cells, and found that metabolism of the non-essential amino acid asparagine (Asn) was highly upregulated. Asn was conditionally essential to B cells, and its synthetic enzyme, asparagine synthetase (ASNS) was upregulated following their activation, particularly more markedly in the absence of Asn, through the integrated stress response sensor general control non-derepressible 2 (GCN2). When *Asns* is deleted B cell survival and proliferation in low Asn conditions were strongly impaired, and removal of environmental Asn by asparaginase or dietary restriction markedly compromised the GC reaction, impairing affinity maturation and the humoral response to influenza infection. Using stable isotope tracing and single cell RNA sequencing, we found that metabolic adaptation to the absence of Asn requires ASNS, and that oxidative phosphorylation, mitochondrial homeostasis, and synthesis of nucleotides was particularly sensitive to Asn deprivation. Altogether, we reveal that Asn metabolism acts as a key regulator of B cell function and GC homeostasis.

**The one sentence summary:** Asparagine metabolism is a critical regulator of B cell function, maintaining the germinal centre reaction.

## Introduction

The germinal centre (GC) reaction is essential for effective humoral immunity^1^. Following encounter with antigen, B cells organise in secondary lymphoid tissue and enter the GC reaction, a cyclic process in which somatic hypermutation (SHM) in a microanatomic region known as the dark zone (DZ) leads to random mutation of immunoglobulin genes. B cells then compete to interact with and receive help from T follicular helper (T_FH_) cells in the microanatomic light zone (LZ). This process continues as affinity maturation occurs and memory B cells or plasma cells are generated.

GC B cells have some of the highest proliferation rates of all mammalian cells, yet their metabolism is unusual and incompletely understood^2^. Unlike most rapidly dividing immune cells, GC B cells predominantly use fatty acid oxidation and oxidative phosphorylation (OxPhos) rather than glycolysis^3–7^, despite residing in a hypoxic and poorly vascularised micro-environment^8^. The key metabolic pathways that are important in their homeostasis have not been fully defined.

Amino acid availability is a critical regulator of T cell responses, demonstrated either by alteration of environmental abundance, synthesis, or interference with cellular import through solute transporters^9–15^. Amino acids are fundamentally required for protein synthesis, but are central to other metabolic processes^16^, and it is apparent in T cells that availability of specific amino acids can have profound effects on their homeostasis.

In contrast, although it is known that B cell-derived lymphomas are sensitive to amino acid deprivation^17–20^, and whilst loss of the capacity to synthesise serine, or deletion of CD98, a component of large neutral amino acid transporters, impairs GC formation, the importance of amino acid availability more broadly in B cell metabolism and humoral immunity is poorly understood^21^.

Here, we find that GC B cells have high levels of amino acid uptake and protein synthesis in vivo. Using integrated multiomic analysis we identify asparagine (Asn), a non-essential amino acid, as a critical regulator of B cell homeostasis and GC maintenance. Mice with conditional deletion of asparagine synthetase (ASNS) in B cells have impairment of GC formation upon depletion of environmental Asn, compromising humoral immunity. ASNS is regulated by sensing through the integrated stress response, and whilst most Asn in B cells is obtained from the extracellular environment following activation, mechanistically, loss of ASNS leads to profound metabolic dysfunction characterised by impairment of OxPhos and failure of nucleotide synthesis, which can be rescued by nucleotide supplementation.

## Results

### GC B cells have highly active protein synthesis and asparagine metabolism

To evaluate protein synthesis rates in GC B cells, we immunised mice with sheep red blood cells (SRBC) and first examined expression of CD98, which forms heterodimers with the amino acid transporter protein SLC7A5 to make L-type amino acid transporter 1 (LAT1), at day 9 following immunisation. As previously reported^21^, CD98 was markedly upregulated in GC B cells compared with IgD^+^ naïve B cells (fig. S1A-B), suggesting high amino acid uptake. To more directly estimate protein synthesis rates in GC B cells *in vivo* and in their spatial context, we used bio-orthogonal non-canonical amino acid tagging (BONCAT), with the amino acid analogues L-azidohomoalanine (L-AHA), and O-propargyl-puromycin (OPP) visualised in situ using Click chemistry^22^. OPP is a puromycin analogue which enters the acceptor site of ribosomes and is incorporated into nascent proteins. L-AHA is an unnatural amino acid, which does not lead to ribosomal stalling once included in polypeptides. To label GC B cells, we immunised mice expressing Rosa26^STOP^tdTomato under the control of *Aicda*-*Cre*, with SRBC. At day 14, we first injected a pulse of L-AHA and then after four hours a pulse of OPP, with analysis one hour later (Fig. 1A). We then used distinct Click chemistry reactions to fluorescently co-label OPP and L-AHA in splenic sections. We found that incorporation of L-AHA and OPP were markedly upregulated in the GC compared with the surrounding follicle of naïve B cells, indicating highly active protein synthesis (Fig. 1B). We also labelled GC B cells with OPP *ex vivo*, and quantified its incorporation by flow cytometry. This confirmed a significant increase in OPP signal in GC B cells compared with naïve IgD^+^ B cells (Fig. 1C-D). Within the GC B cell population, signal was highest in dark zone (CXCR4^hi^ CD86^low^) GC B cells, which are undergoing proliferation and SHM (Fig. 1C-D). GC B cells therefore have high rates of both amino acid transporter expression and protein synthesis.

**Figure 1.**
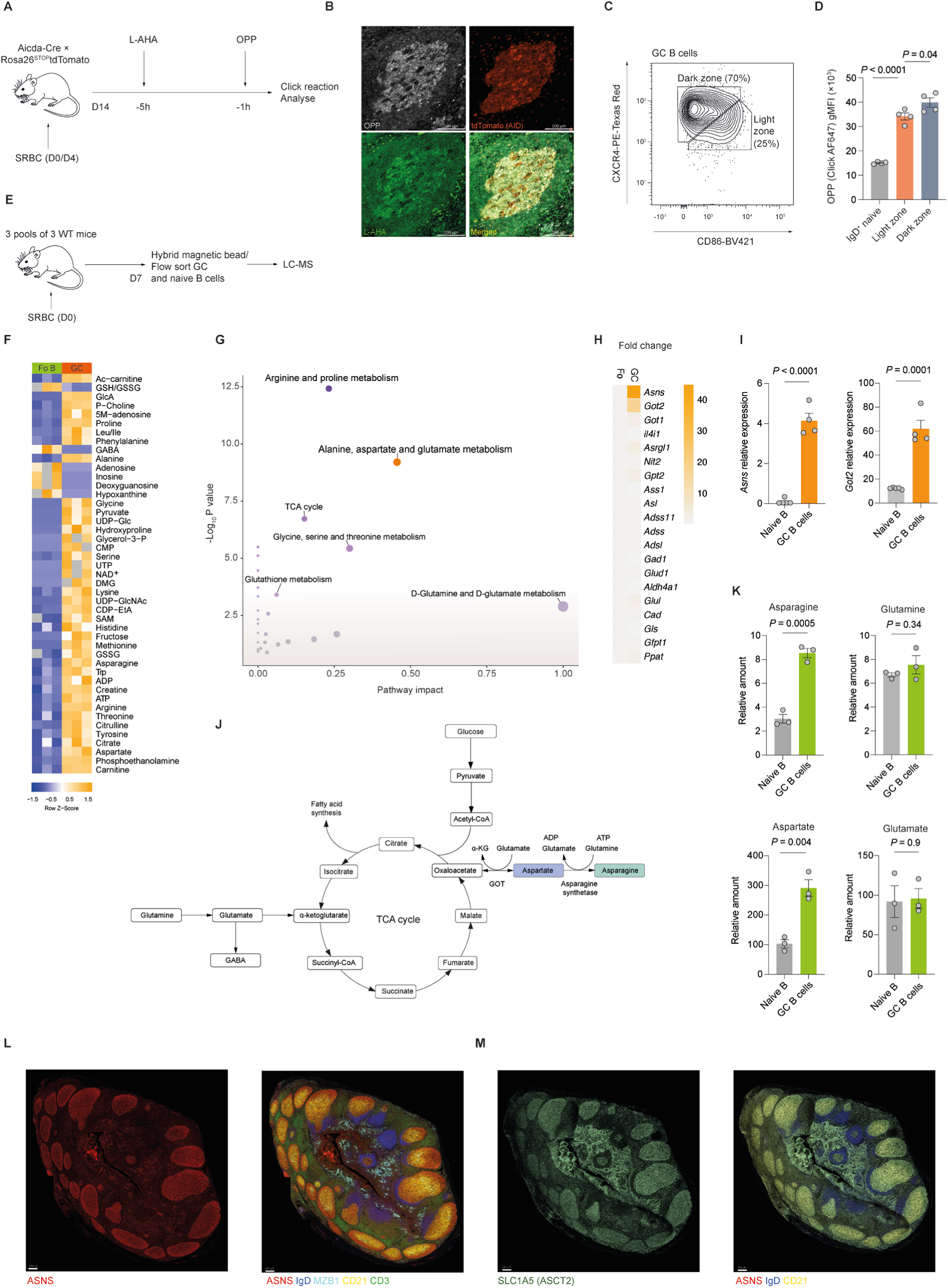
GC B cells have highly active protein synthesis and asparagine metabolism. **A.** Schematic of in vivo bio-orthogonal non-canonical amino acid tagging. Aicda-cre × Rosa26^STOP^tdTomato mice were immunised with SRBC and then boosted at day 4. At day 14, they were injected with L-AHA, and then four hours later with OPP. They were sacrificed one hour later, and non-canonical amino acid incorporation into splenic tissue was revealed using Click chemistry. **B.** Representative immunofluorescence images of splenic GCs labelled with OPP, tdTomato (AID), and L-AHA. Representative of two independent experiments. Scale bar: 100μm. **C.** Gating strategy of light zone (CD86^hi^CXCR4^lo^) and dark zone (CD86^lo^CXCR4^hi^) GC B cells **D.** Quantification of OPP incorporation in naïve, LZ GC and DZ GC B cells from n=4 mice. Representative of two independent experiments. **E.** Schematic of experimental design. Three pools of three WT mice were immunised with SRBC, and at day 7 GC and follicular B cells were isolated by a hybrid magnetic bead and flow sorting approach, before analysis of polar metabolites by LC-mass spectrometry (LC-MS). **F.** Heatmap of significantly differentially abundant metabolites (P_adj_ <0.05) in naïve and GC B cells at day 7 from wild type mice immunised with SRBC, measured by LC-MS (n=3 pools of 3 mice/pool). Representative of two independent experiments. **G.** Integrated pathway analysis of differentially abundant metabolites from **F** and expressed genes from ImmGen in GC compared to naïve B cells (GSE15907). **H.** Heatmap of relative gene expression of KEGG Alanine, Aspartate, and Glutamate pathway in GC vs naïve B cells, represented as fold change. Data are from GSE133971^25^. **I.** Relative expression of *Asns* and *Got2* in flow-sorted GC and naïve B cells from wild type mice immunised with SRBC at day 7, normalised to *Ubiquitin C* (*Ubc*) (n=4 mice). Representative of two independent experiments. **J.** Diagram of asparagine synthesis and its association with the TCA cycle. **K.** Relative amounts of asparagine, aspartate, glutamine, and glutamate in GC and naïve B cells measured by LC-MS (n=3 pools of 3 mice) as in **F**. Representative of two independent experiments. **L.** Multiplexed CellDIVE images of normal human tonsil section, with immunofluorescence staining for ASNS, CD21, IgD, MZB1 and CD3. Representative of 3 separate tonsils. Scale bar = 250μm. **M.** Multiplexed CellDIVE images of normal human tonsil serial section of **L**, with immunofluorescence staining for SLC1A5 (ASCT2), CD21 and IgD. Representative of 3 separate tonsils. Scale bar = 250μm. Statistical significance was determined by two-way ANOVA with Tukey’s multiple testing correction (F), hypergeometric test (G), unpaired two-tailed t test (I,K) or one-way ANOVA with Tukey’s multiple testing correction (D). Data are presented as the mean +/− SEM.

We next directly profiled the metabolite content of GC B cells. To do so, we isolated GC and naïve B cells from pools of wild type mice following immunisation with SRBC and quantified key metabolites using liquid chromatography-mass spectrometry (LC-MS) (Fig. 1E). This revealed a broad increase in the levels of metabolites in GC B cells, with overrepresentation of intermediates of glycolysis, the TCA cycle, and most amino acids (Fig. 1F). Notably however, nucleotide precursor molecules (e.g.adenosine and hypoxanthine) were depleted in GC B cells, potentially reflective of their consumption during the nucleic acid synthesis required for high rates of proliferation. To refine our approach, we fused metabolite and transcriptional datasets^23^ and performed integrated pathway analysis on the combined data using hypergeometric set testing^24^ (Fig. 1G). We found that there was enrichment of amino acid metabolic pathways in GC B cells, most notably the Kyoto Encyclopedia of Genes and Genomes (KEGG) Alanine, Aspartate, and Glutamate (AAG) metabolism pathway. Examination of the leading-edge genes of the AAG pathway in an independent transcriptional dataset revealed an approximately 40-fold increase in the expression of asparagine synthetase (*Asns*) and a 15-fold increase in glutamic oxaloacetic transaminase-2 (*Got2*) in GC B cells compared with naïve follicular B cells^25^ (Fig. 1H). We confirmed upregulation of *Asns* and *Got2* in sorted GC B cells by qPCR (Fig. 1I) and by immunoblotting (fig. S1C).

ASNS synthesises Asn from aspartate (Asp) in an ATP-dependent reaction, and GOT2 synthesises Asp from the TCA cycle intermediate oxaloacetate^26^ (Fig. 1J). The relative levels of both Asn and Asp were markedly increased in GC B cells compared with naïve B cells, which had equivalent glutamate (Glu) and glutamine (Gln) levels (Fig. 1K).

To examine ASNS expression in its spatial context, we performed multiplex imaging of human tonsils (Fig. 1L and fig. S1D). ASNS was strongly expressed in GCs and MZB1^+^ plasmablasts. There was negligible expression in the IgD^+^ B cell follicle or in the CD3^+^ T cell zone (fig. S1E). ASNS levels were spatially regulated within the GC, with higher expression in the DZ, where proliferation and SHM occurs (fig. S1D-E). We also performed immunostaining for the amino acid transporter Alanine serine cysteine transporter 2 (ASCT2), the main transporter of Asn and which also transports L-AHA^13,27^, and found high expression in GC B cells, more pronounced in the DZ like ASNS (Fig. 1M and fig. S1F). Asn metabolism is therefore one of the most upregulated pathways in GC B cells and plasmablasts, suggesting that it may play an important role in their physiology.

### B cells require ASNS when the availability of Asn is limited

To approximate the concentration of Asn and Gln encountered by B cells in secondary lymphoid tissue, we eluted interstitial fluid from pooled lymph nodes (LNIF) of wild type mice, and compared this to serum. We found that Asn levels in LNIF were substantially higher than those in serum, whilst Gln levels were comparable (Fig. 2A). This suggested that lymphocytes may be adapted to the higher local Asn concentration in their niches relative to serum levels.

**Figure 2.**
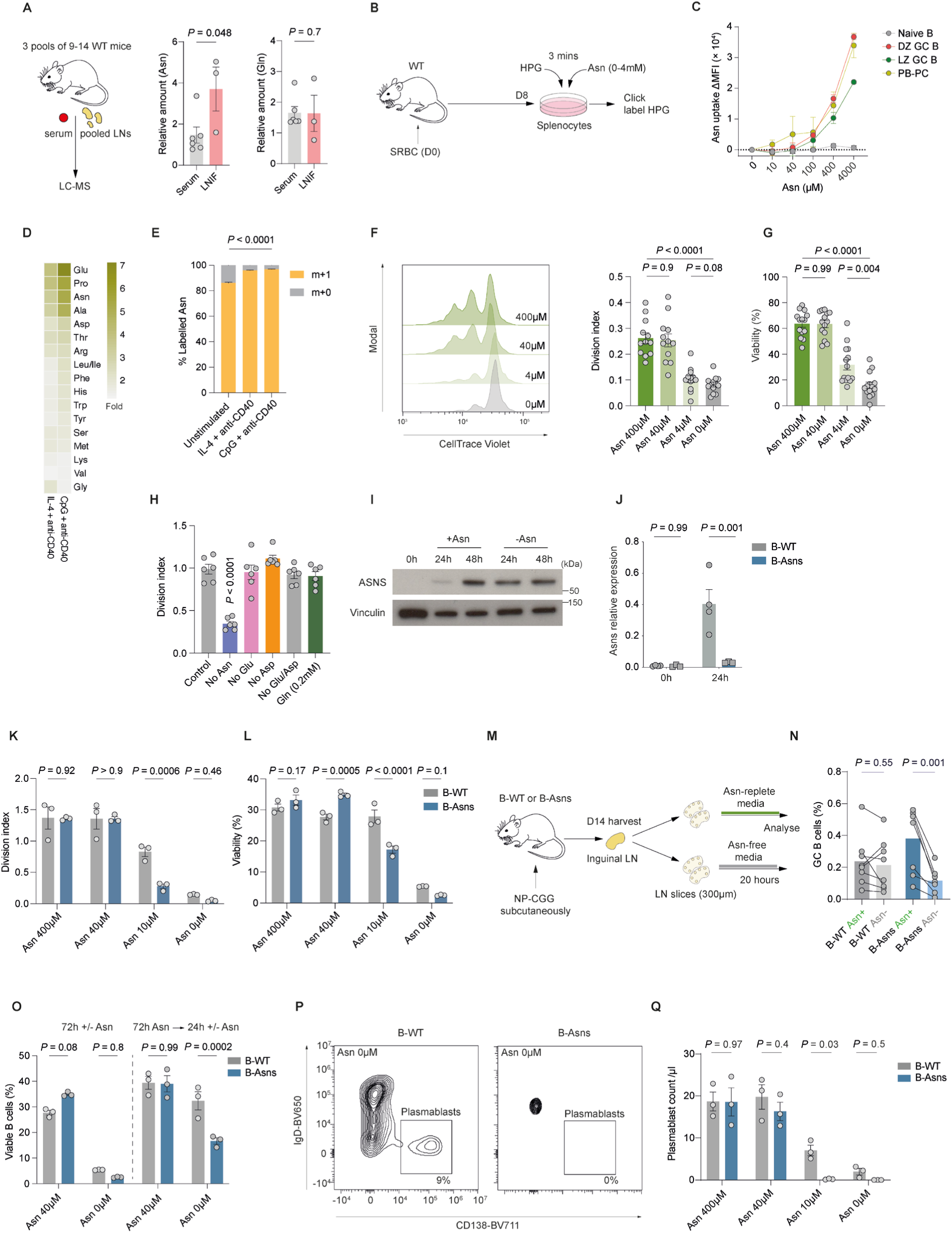
B cells require ASNS when the availability of Asn is limited. **A.** Schematic of serum and pooled lymph node (axial, brachial, inguinal, cervical, mediastinal, mesenteric) collection and LC-MS strategy and quantification of relative amounts of asparagine (Asn) and qlutamine (Gln) in serum and lymph node intersistial fluid measured by LC-MS (n=3 pools of 9-14 unimmunised WT mice for LNIF and 2-3 unimmunised WT mice for serum). Pooled from three independent experiments. **B.** Schematic of ex vivo ASCT2 (SLC1A5)-mediated competitive Asn uptake assay. **C.** Quantification of the shift in the HPG signal intensity (ΔMFI) as in **fig. S2A** reflecting Asn uptake calculated as following: Asn uptake at *x* concentration = HPG MFI_[Asn0]_ − HPG MFI_[Asn *x*]_. Splenic IgD^+^GL-7^−^ naïve B, IgD^−^CD138^+^ plasmablast/plasma cells (PB-PC), CD38^−^GL-7^+^CXCR4^lo^CD86^hi^ LZ and CD38^−^GL-7^+^CXCR4^hi^CD86^lo^ DZ GC B cells harvested from n=4 mice at day 8 following SRBC immunisation. Background signal subtraction was performed based on 4°C and no-HPG added controls. Representative of two independent experiments. **D.** Heatmap of fold change of indicated amino acids in B cells, measured by LC-MS following stimulation for 24h with IL-4 and anti-CD40, or CpG and anti-CD40, relative to unstimulated condition (n=3 mice). Representative of two independent experiments. **E.** Fraction of labelling of free intracellular Asn with ^15^N in B cells stimulated as in **D**, in the presence of ^15^N_1_-Asn (n=3 mice). Representative of two independent experiments. **F.** Representative histogram of dilution of CellTrace Violet in B cells stimulated with IL-4 and anti-CD40 and cultured for 72h with the indicated concentration of Asn. **G.** Division index and viability of B cells stimulated with IL-4 and anti-CD40 and cultured for 72h with the indicated concentration of Asn. Each dot represents a single mouse. Pooled from three independent experiments. **H.** Division index of B cells stimulated for 72h with IL-4 and anti-CD40 in the presence or absence of the indicated amino acids (n=6 mice). Data pooled from three independent experiments. **I.** Representative immunoblot of ASNS in B cells stimulated with IL-4 and anti-CD40 with or without Asn, at the indicated timepoints. Representative of two independent experiments. **J.** Relative expression of *Asns* by qPCR in freshly isolated, and stimulated (IL-4 and anti-CD40 for 24h) B cells (n=4 B-WT and n=4 B-Asns mice). Representative of two independent experiments. **K.** Division index of B cells from B-Asns and B-WT mice stimulated for 72h with IL-4 and anti-CD40 at the indicated concentration of Asn (n=3 B-WT and n=3 B-Asns mice). Data pooled from two independent experiments and representative of >3 independent experiments. **L.** Viability of B cells from B-Asns and B-WT mice stimulated for 72h with IL-4 and anti-CD40 at the indicated concentration of Asn (n=3 B-WT and n=3 B-Asns mice). Data pooled from two independent experiments and representative of >3 independent experiments. **M.** Schematic of lymph node slice culture system. B-WT and B-Asns mice were immunised with NP-CGG in alum subcutaneously and inguinal lymph nodes were harvested at day 14. 300μm thick slices were cultured for 20h in the presence or absence of Asn in duplicates and analysed by flow cytometry. **N.** Quantification of CD38^−^GL-7^+^ GC B cell proportions by flow cytometry in lymph node slices cultured for 20h in the presence or absence of Asn (400μM). Each dot represents single lymph node (n=8 lymph nodes from n=4 mice). Pooled from two independent experiments. **O. Left panel:** Viability of B cells from B-Asns and B-WT mice stimulated for 72h with IL-4 and anti-CD40 at the indicated concentration of Asn (40μM), data from **L**. **Right panel:** Viability of B cells stimulated with IL-4 and anti-CD40 with 40μM Asn for 72h, then restimulated for an additional 24h in the presence (40μM) or absence of Asn (n=3 mice). Data pooled from two independent experiments. **P.** Representative flow cytometry plots of IgD^−^CD138^+^ plasmablasts generated following stimulation of B cells from B-WT and B-Asns mice with LPS and IL-4 for 72h at the indicated concentration of Asn. **Q.** Quantification of plasmablast counts per µl as in **P**. Data pooled from three independent experiments. Statistical significance was determined by a two-tailed Mann–Whitney U-test (A), repeated measures one way ANOVA with Tukey’s multiple testing correction (F-G) and ordinary one way ANOVA with Dunnet’s multiple testing correction (E,H), paired two-tailed t test (N) or two-way ANOVA with Šidák’s multiple testing correction (J-L,O,Q). Data are presented as the mean +/− SEM.

Asn is a non-essential amino acid and can be either acquired exogenously or synthesised by ASNS. To understand these relative contributions to Asn metabolism, we first functionally evaluated Asn uptake in *ex vivo* GC B cells, using an indirect assay in which the bio-orthogonal amino acid analogue homopropargylglycine (HPG) competes for transport through ASCT2 with Asn (Fig. 2B). HPG is then fluorescently labelled using Click chemistry, and its uptake measured by flow cytometry^27^. We found that there was clear dose-dependent competition between HPG and Asn in GC B cells and plasmablasts (fig. S2A). This was most marked in DZ GC B cells, in keeping with their high levels of OPP incorporation and ASNS expression, and indicating substantially elevated Asn uptake compared to naïve B cells (Fig. 2C).

To understand the dynamics of Asn uptake following B cell activation, we then stimulated wild type B cells in vitro with either the TLR9 agonist CpG and agonistic anti-CD40 antibody, or IL-4 and agonistic anti-CD40, for 24h in the presence of ^15^N-labelled Asn, and quantified amino acids by mass spectrometry. We found a generalised increase in the intracellular concentrations of most amino acids, but in particular proline (Pro), glutamate (Glu), and Asn (Fig. 2D). The majority of Asn was labelled with ^15^N in all conditions, indicating high exogenous uptake, and in preference to endogenous synthesis, which is bioenergetically demanding (Fig. 2E).

Next, to determine the requirements of B cells for exogenous Asn, we stimulated them with agonistic anti-CD40 and IL-4 in RPMI-1640 media with varying concentrations of Asn. Standard RPMI-1640 contains higher concentrations of Asn (378μM) compared with those found in plasma (~40μM) and cerebrospinal fluid (CSF) (~4μM)^28,29^. When B cells were deprived of Asn or supplemented with a low concentration of Asn (4μM), and stimulated with IL-4 and anti-CD40 for 72h, there was a severe reduction in cell proliferation and viability, compared with supplementation of Asn at 40μM or 400μM concentrations (Fig. 2F-G). We noted no decrease in cell proliferation or viability on withdrawal of exogenous aspartate (Asp) and/or Glu (Fig. 2H and fig. S2B). There was a reduction in the expression of the activation markers CD86 and MHCII following Asn deprivation, nascent protein synthesis measured by OPP incorporation was lower at 24h (fig. S2C-E), and apoptosis was increased (fig. S2F). ASNS levels in B cells increased substantially over time following stimulation, and the absence of Asn led to higher ASNS expression at 24h (Fig. 2I). ASNS was increased with synergistic IL-4 and anti-CD40 stimulation (fig. S2G).

To determine the role of Asn synthesis in B cell homeostasis, we generated *Cd79a*-*Cre* × *Asns* ^LoxP^ mice (B-Asns hereafter), in which Cre recombinase is expressed at earliest stages of B cell development^30^. There was no evident B cell developmental defect and the mature B cell compartment was normal, with effective deletion of *Asns*. (Fig. 2J, fig. S2H-J). Survival and proliferation were severely compromised in B-WT B cells, and to a greater extent in B-Asns B cells, when Asn was restricted in the culture medium, but were normal under Asn-replete conditions (40µM and 400µM) (Fig. 2K-L). *In vitro* culture systems lack the spatial organisation and cellular complexity of lymphoid tissue, and may therefore reproduce normal immune dynamics less well. We used a live *ex vivo* lymph node slice platform to investigate how Asn deprivation affected GC B cells from mice immunised with 4-hydroxy-3-nitrophenylacetyl-chicken gamma globulin (NP-CGG) in alum (Fig. 2M)^31^. We found that GC B cells from B-Asns lymph nodes were much reduced after 20h of culture without Asn, but total B cell proportions were unaffected (Fig. 2N, fig. S2K). Since GC B cells express ASNS, and B cells upregulate ASNS following stimulation, we reasoned that it conferred protection against Asn deprivation. We therefore pre-stimulated B-Asns or B-WT B cells in the presence of Asn, which was then withdrawn for 24 hours. In contrast to when Asn was initially absent, B-WT B cells were minimally affected by the absence of Asn (Fig. 2O). However, in B-Asns B cells this protective effect was attenuated (Fig. 2O). Plasmablast differentiation induced by LPS and IL-4 was largely abolished in B-Asns B cells when Asn was absent or at low concentrations (10μM), but remained unchanged in the presence of Asn (Fig. 2P-Q, fig. S2L).

Asn uptake and synthesis is therefore specifically required during initial B cell activation, and loss of synthetic capacity leads to severely compromised cell division and survival in B cells in low Asn conditions.

### The GC reaction is sensitive to Asn deprivation

To understand the effect of Asn deprivation on the GC reaction *in vivo*, we treated B-Asns and B-WT mice with asparaginase (ASNase), an enzyme of bacterial origin which rapidly and effectively hydrolyses Asn and is a well-established treatment for leukaemia^32^. We used two treatment schedules (Fig. 3A). In the ‘standard’ schedule, we administered ASNase from one day prior to immunisation with SRBC, and then every two days over a nine day period before analysis. This includes the period of GC establishment and expansion. In the ‘post-formation’ schedule, we used only two doses, at days six and eight following immunisation, therefore targeting established GCs. ASNase effectively depleted Asn without affecting Gln in the serum, and minimally affected naïve B cells, marginal zone B cells, and T_FH_ and T_FR_ T cells (fig. S3A-G).

**Figure 3:**
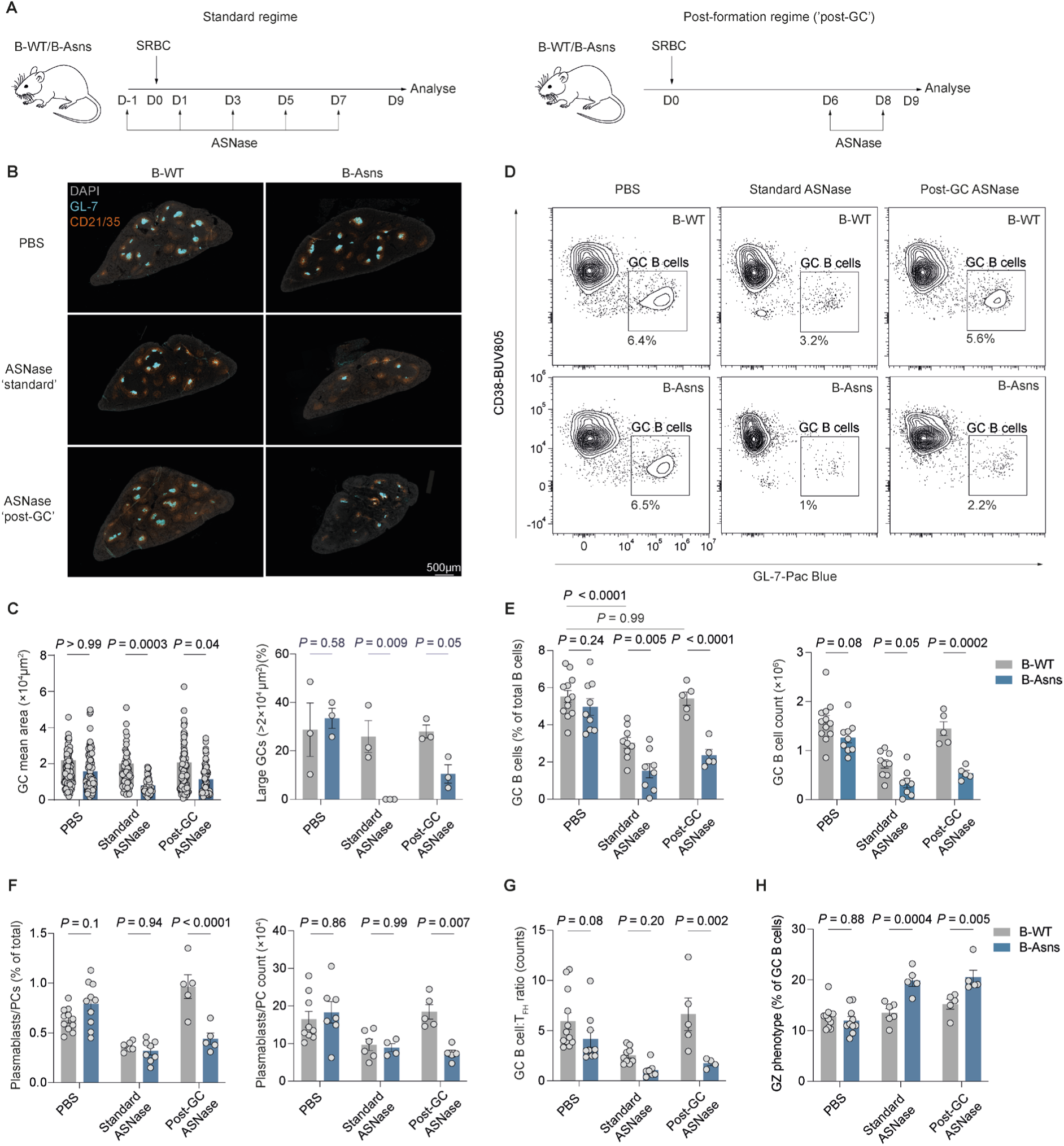
The GC reaction is sensitive to Asn deprivation. **A.** Schematic of *in vivo* ASNase administration regimes. Standard regime: B-WT or B-Asns mice were intraperitoneally injected with ASNase (500U/kg: ≈10U/mouse) or PBS vehicle one day prior to immunisation with SRBC (day 0) and given repeated ASNase injections every two days until day 7. Spleens were harvested at day 9 and analysed by immunohistochemistry and flow cytometry. Post-formation ‘post-GC’ regime: B-WT or B-Asns mice were intraperitoneally injected with SRBC (day 0) and given two ASNase (500U/kg: ≈10U/mouse) or PBS injections at days 6 and 8 with analysis at day 9. **B.** Immunohistochemistry of representative spleen sections (8μm) at day 9 post SRBC immunisation. GL-7 and CD21/35 highlights GCs and B cell follicles, respectively. DAPI is used as background staining. Scale bar 500μm. Data pooled from three independent experiments. **C.** Image quantification of splenic GCs from **B**. **Left panel**: mean area of individual GCs (μm^2^). Each data point represent a GC pooled from n=3 mice per condition. **Right panel**: frequency of GCs larger than 20000µm^2^ (0.02mm^2^). Each data point indicates a mouse. n=3 from each condition. Data pooled from three independent experiments **D.** Representative flow cytometry plot of GC B cells (CD19^+^CD38^−^GL-7^+^) at day 9 post-immunisation with SRBC, from B-WT and B-Asns mice treated with PBS, standard ASNase or post-GC ASNase. **E.** Quantification of splenic CD38^−^GL-7^+^ GC B cell proportions (% of CD4^−^CD19^+^ B cells) and absolute counts. Each data point represents a single mouse. Data pooled from >3 independent experiments. **F.** Quantification of IgD^−^IRF4^+^CD138^+^ splenic plasmablast/plasma cell proportions (% of total) and absolute counts. Each data point represents a single mouse. Data pooled from >3 independent experiments. **G.** Quantification of the GC B cell numbers relative to T_FH_ numbers gated as CXCR5^hi^ PD-1^+^ within CD19^−^ CD4^+^ FoxP3^−^ T cells. Each data point represents a single mouse. Data pooled from >3 independent experiments. **H.** Quantification of the proportion of the grey zone (GZ) (CD86^hi^CXCR4^hi^) GC B cells (as % of GC B cells). Data pooled from >3 independent experiments. Statistical significance was determined by two-way ANOVA with Šidák’s multiple testing correction. Data are presented as the mean +/− SEM.

We found that following ASNase administration, the GC reaction was markedly compromised in B-Asns mice with both regimes, with reduced GC B cell and plasmablast proportions, numbers, and GC area (Fig. 3B-F, fig. S3H-I). B-WT mice however, were insensitive to the post-formation regime. The effect of genotype on plasmablast numbers was only evident with later ASNase treatment. There was a severe reduction in GC B cells relative to T_FH_ numbers in B-Asns mice treated with the post-GC formation regime, suggesting a selective impact on GC B cells (Fig. 3G).

We then measured the effect of post-GC formation ASNase on the humoral response to the protein-hapten conjugate immunogen NP-CGG (fig. S4A-B). This ASNase regime reduced the levels of both low- and high-affinity antibodies (NP_20_ and NP_2_, respectively), with minimal genotype effect (fig. S4C).

Within the GC B cell compartment, there was a reduction in DZ phenotype cells in treated B-Asns mice, but an increase in those with a ‘grey zone’ (GZ) phenotype (Fig. 3H and fig. S4D-F). The GZ is the compartment of peak cell proliferation in the GC, and is characterised by high expression of the cell cycle protein cyclin B1, and of cells in G2/M phase^33^(fig. S4G). We did not detect a defect in S phase in GC B cells, as measured by incorporation of the thymidine analogue EdU *in vivo*, which was slightly increased following ASNase treatment (fig. S4H-I).

Deprivation of Asn with ASNase therefore strongly compromises the GC reaction in the absence of ASNS and possibly as a consequence, plasmablast formation.

### Asn is required for GC B cell function

In order to examine the requirement for Asn in a more physiologically relevant setting, and over a longer time course, we switched to dietary limitation of Asn. To test the cell intrinsic effect of *Asns* deletion, and to minimise confounding due to cage effects or food intake, we first generated mixed chimeras with CD45.1 wild type and CD45.2 B-Asns or B-WT bone marrow. After 8 weeks, mice were given either an Asn-free or normal diet for 12 days, immunised with SRBC, and analysed after a further 9 days of dietary modification (fig. S5A).

The relative proportion of GC B cells in CD45.2 B-Asns B cells was much lower in mice fed an Asn-free diet, and the DZ/LZ ratio was reversed compared to B-WT cells (fig. S5B-C), which reproduced our findings with ASNase. Also reduced were splenic plasmablasts of B-Asns origin (fig. S5D). We found a marked reduction in the frequency of SRBC-binding B-Asns-derived GC B cells in mice fed an Asn-free diet (fig. S5E-F), suggesting impaired development of antigen-binding clones.

We then examined the effect of dietary Asn deprivation on the humoral response to NP-CGG in a non-chimeric setting, in B-Asns and B-WT mice (Fig. 4A). This allowed us to follow the GC response over time by measuring antibody affinity maturation.

**Figure 4:**
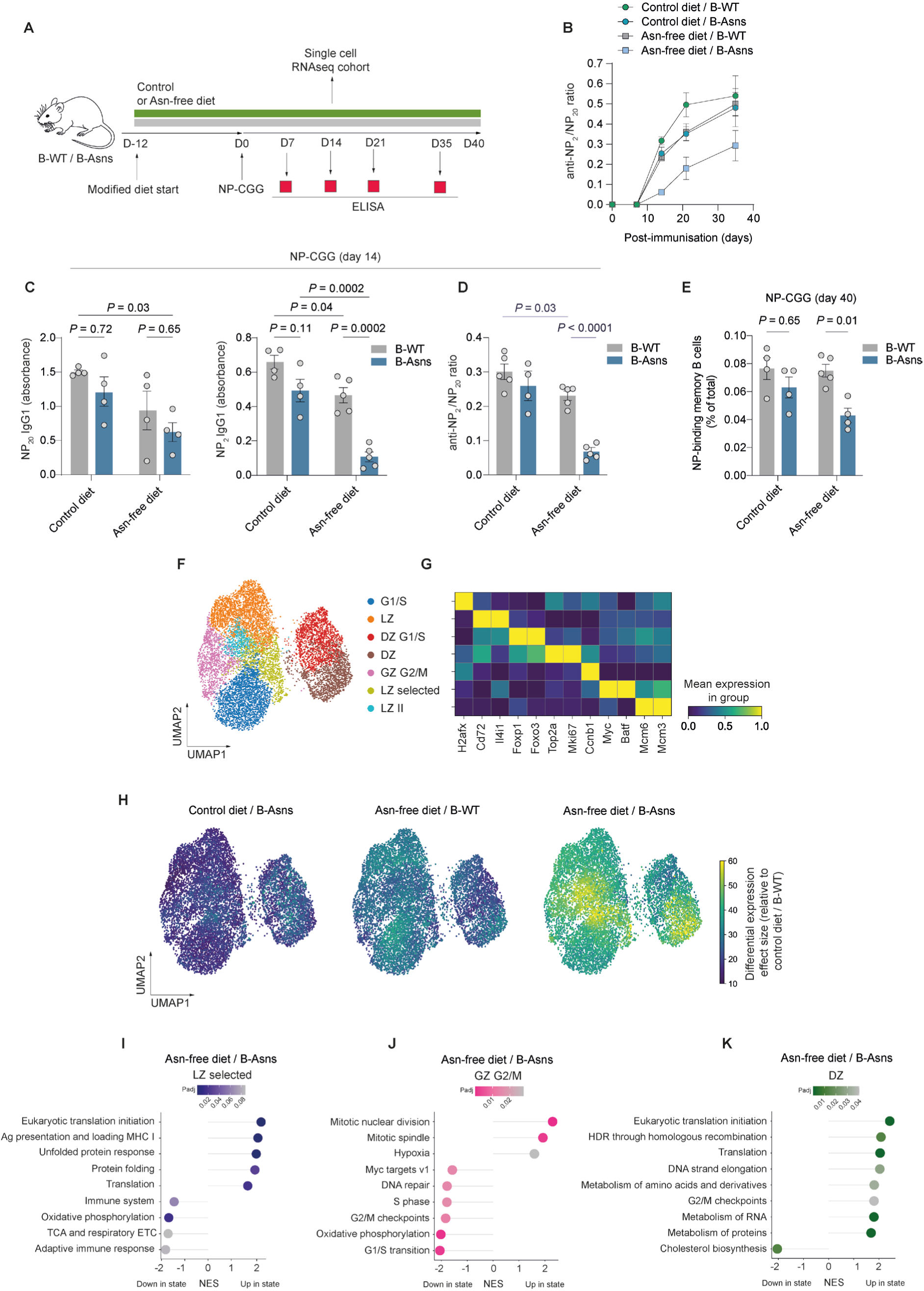
Asn is required for GC B cell function. **A.** Schematic of non-chimeric NP-CGG/dietary modification experiment. B-WT or B-Asns mice were assigned to either an Asn-free or control diet (Asn-replete) for 12 days, before immunisation with 50μg NP-CGG precipitated in Alum hydrogel (1:1) subcutaneously on both flanks. Mice were bled on day 7, 14, 21 and 35 post-immunisation and spleens were analysed following sacrifice at day 40. A separate experimental cohort undergoing similar dietary modification regime was immunised intraperitoneally with 100μg NP-CGG and sacrificed on day 14 for scRNAseq analysis of splenic GC B cells. **B.** Comparison of the ratio of IgG1 NP-specific high-affinity antibodies to low-affinity antibodies detected by binding to NP_2_ and NP_>20_ antigens, respectively, from B-WT/control diet (n = 4), B-Asns/control diet (n = 4), B-WT/Asn-free diet (n = 5), B-Asns/Asn-free diet (n=5) across different time points. Data pooled from two independent experiments. **C.** Quantification of IgG1 anti-NP antibodies (NP_>20_-BSA at 1:1600 and NP_2_-BSA at 1:200 dilution respectively) at day 14, n=4-5 mice each condition. Data pooled from two independent experiments. **D.** Quantification of NP_2_:NP_20_ ratio (at 1:200 dilution) of IgG1 anti-NP antibodies at day 14, n=4-5 mice each condition. Data pooled from two independent experiments. **E.** Comparison of the proportions of splenic NP-binding memory B cells gated as CD19^+^ B220^+^IgD^−^GL-7^−^CD138^−^CD38^+^ within total cells at day 40. Data pooled from two independent experiments. **F.** UMAP and cluster annotation based on multiresolution variational inference (MrVI) latent space of integrated control diet/B-WT (n=2550 cells), control diet/B-Asns (n=2971 cells), Asn-free diet/B-WT (n=1944 cells), and Asn-free diet/B-Asns (n=2700 cells) (n = 3 mice per condition), as in **A**. **G.** Heatmap of selected differentially expressed genes used to identify clusters, as in **F**. **H.** MrVI cluster-free differential gene expression effect size for the indicated conditions relative to the control diet/B-WT group. **I.** GSEA of the indicated pathways in the ‘LZ selected’ cluster of Asn-free diet/B-Asns relative to control diet/B-WT groups. **J.** GSEA of the indicated pathways in the in the ‘GZ’ cluster of Asn-free diet/B-Asns relative to control diet/B-WT groups. **K.** GSEA of the indicated pathways in the in the ‘DZ’ cluster of Asn-free diet/B-Asns relative to control diet/B-WT groups. Statistical significance was determined by two-way ANOVA with Tukey’s multiple testing correction (C-D) or Šidák’s multiple testing correction (E) and adaptive multi-level split Monte-Carlo scheme (I-K). Data are presented as the mean +/− SEM.

At day 14, levels of IgG1 anti-NP antibodies binding to high conjugation NP-BSA (NP_20_), were lower in B-Asns mice deprived of Asn, and there was substantially reduced binding to low conjugation NP_2_ in sera, also seen in B-WT mice given an Asn-free diet (Fig. 4B-D, fig. S5G), which was maintained over time. This indicated that affinity maturation was defective in this setting. We did not observe a large difference in anti-NP IgM, which is of likely extrafollicular origin (fig. S5H). As a more refined approach for studying the extrafollicular response, we also examined the anti-NP IgM antibody response at day 7 after immunisation with the T-independent antigen NP-Ficoll^34^ (fig. S5I). There was a reduction in anti-NP_20_ antibody levels in B-Asns mice on an Asn-free diet, compared to B-WT given a control diet (fig. S5J).

Forty days following immunisation with NP-CGG, we found that there was a substantial decrease in the NP-binding memory B cell compartment (Fig. 4E, fig. S6A). We also performed experiments in which mice were immunised with NP-CCG a week prior to dietary modification, and then at day 36 boosted with further NP-CGG (fig. S6B-C). This showed attenuation of the boost-associated increase in high-affinity anti-NP antibody levels at day 49 in B-Asns mice given an Asn-free diet (fig. S6D-E), while serum Asn levels remained unchanged around this time, likely due to systemic compensation (fig. S6F).

Next, we gave B-WT and B-Asns mice a control or Asn-free diet for 12 days, immunised them with NP-CGG, then 14 days later sorted GC B cells and performed single cell RNA sequencing (as depicted in Fig. 4A). We trained a multiresolution variational inference (MrVI) model and then clustered cells using its *u* latent space^35^. MrVI generates two separate latent spaces; one based on all cells, and another on individual samples, therefore allowing annotation-free comparative analysis. We distinguished 7 clusters, with varying light zone, dark zone, and grey zone marker gene expression (Fig. 4F-G). Examining differential gene expression across all cells with MrVI, we found a gradient across conditions relative to B-WT mice on control diet, with the largest effect size in B-Asns mice given an Asn-free diet (Fig. 4H). When cell state differential abundance was analysed, there were differences in all conditions compared to B-WT mice on control diet, centred on the dark zone (fig. S6G).

We next performed gene set enrichment analysis (GSEA), using a conventional pseudobulk differential gene expression approach. Focusing on B-Asns mice given an Asn-free diet and examining clusters which showed the largest differential gene expression effect, we found that in the LZ-selected cluster, which expresses markers of GC B cell selection such as *Myc* and *Batf*, there was downregulation of OxPhos and TCA cycle sets, and upregulation of unfolded protein response gene sets (Fig. 4I). We next examined the GZ G2/M cluster, given our findings in B-Asns mice treated with ASNase (Fig. 4J). We found increased expression of mitosis gene sets, and downregulation of G2/M checkpoint, G1/S transition, and S phase gene sets, as well as OxPhos. In the DZ cluster there was upregulation of cell cycle-related gene sets and downregulation of cholesterol synthesis (Fig. 4K). Examining data from the mixed bone marrow chimera experiment (fig. S5A), we identified defective mitosis within the dark zone of B-Asns GC B cells under dietary Asn restriction (fig. S6H-I).

Having shown that Asn deprivation leads to compromised GC output, we then asked if supplementation with exogenous Asn would enhance it. We administered Asn in drinking water to wild type mice fed a standard diet for 14 days before immunisation with NP-CGG, and 14 days later quantified IgG1 anti-NP antibodies. We did not observe an increase in antibody concentration or affinity maturation (fig. S6J-K). This suggested that the amount of Asn provided by a standard diet was fully sufficient for the GC reaction over this timescale.

Asn synthesis in B cells by ASNS therefore acts to maintain GC B cell function in the face of dietary Asn deprivation.

### Asn metabolism controls the humoral response to influenza infection

The GC reaction is integral to humoral immune responses following flu infection. Therefore, we investigated whether Asn metabolism could affect influenza-induced GC and antibody generation. Mice were assigned to either an Asn-free or normal diet for 12 days, as previously described, and then intranasally infected with the X31 H3N2 influenza A strain (Fig. 5A). Starting from day 3, characteristic weight loss was observed, peaking around day 7 (fig. S7A). Viral clearance during primary influenza infection is largely mediated by the activity of T cells and other non-B cell immune populations, which coincides with the initiation of recovery in mice^36^. We found that the rate of weight recovery under all conditions remained comparable, suggesting that there was no important defect in the non-B cell compartment of experimental mice (fig. S7A).

**Figure 5.**
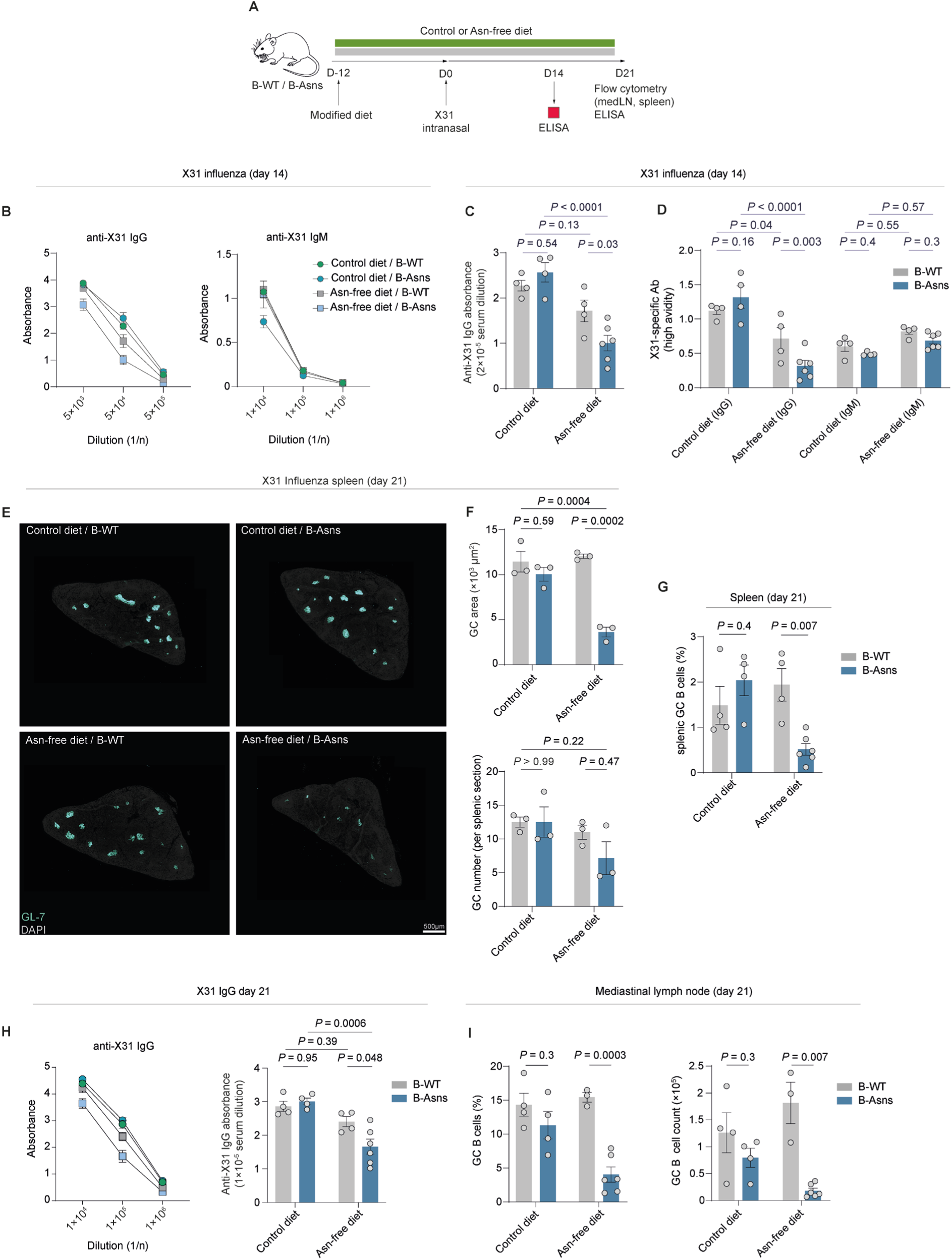
Asn metabolism controls the humoral response to influenza infection. **A.** Schematic of influenza/dietary modification experiment. B-WT and B-Asns mice were transferred to a control (Asn-replete) or Asn-deficient diet for 12 days, before intranasal infection with X31 influenza A strain. Mice were weighed daily for 14 days and bled at days 14 and 21. Mediastinal lymph node and spleen were harvested for flow cytometry and immunohistochemistry analyses following sacrifice at day 21. **B.** Dilution curves of X31-specific IgG and IgM antibodies at day 14. n=4-6 mice each condition. Data pooled from two independent experiments. **C.** Comparison of X31-specific IgG (at 1/5×10^4^ dilution) at day 14. n=4-6 mice each condition. Data pooled from two independent experiments. **D.** Comparison of high-avidity X31-specific IgG and IgM (at 1/5×10^4^ dilution) quantified by values in **C and fig S7B** with matching avidity values in **fig. S7D-E** at day 14. n=4-6 mice each condition. Data pooled from two independent experiments. **E.** Immunohistochemistry of representative spleen sections (8μm) at day 21 post X31 influenza infection. GL-7 highlights GCs. DAPI is used as background staining. Scale bar 500µm. Data pooled from two independent experiments **F.** Image quantification of X31-induced splenic GCs from **E**. Mean GC area (μm^2^) per mouse and GC count per splenic section (average of two non-serially sliced) per mouse. Each data point indicates a mouse. n=3 from each condition. Data pooled from two independent **G.** Flow cytometric quantification of splenic GC B cell (as gated in **fig. S7G**) proportions at day 21. n=4-6 mice each condition Data pooled from two independent experiments. **H.** Dilution curves and quantification of X31-specific IgG antibodies at day 21. n=4-6 mice each condition as in **B**. Data pooled from two independent experiments. **I.** Quantification of GC B cell ratio and counts in mediastinal lymph nodes at day 21. n=3-6 mice each condition. Data pooled from two independent experiments. Statistical significance was determined by two-way ANOVA with Tukey’s multiple testing correction (C,D,F,H) or Šidák’s multiple testing correction (G,I). Data are presented as the mean +/− SEM.

We initially examined the antibody response to X31 influenza in mouse sera. At day 14 post-infection, levels of anti-X31 IgG antibodies were substantially lower in B-Asns mice fed an Asn-free diet (Fig. 5B-C). However, we did not observe a significant difference in anti-X31 IgM levels (Fig. 5B, fig. S7B). Affinity maturation in GCs is essential for the development of influenza specific high-affinity antibodies. We therefore assessed the functional affinity (avidity) of X31-specific antibodies by ELISA, using the chaotropic agent urea at increasing concentrations to elute weakly binding antibodies^37^. We observed that anti-X31 IgG antibodies from B-Asns mice fed an Asn-free diet were significantly more sensitive to urea treatment, suggesting markedly impaired affinity (Fig. 5D, fig. S7C-D). As expected, anti-X31 IgM antibodies exhibited greater sensitivity to urea, to a similar extent across all experimental groups, indicating lower avidity compared to anti-X31 IgG antibodies, owing to the lack of affinity maturation in their extrafollicular origin (Fig. 5D, fig. S7E). Influenza infection induces GCs in the spleen and mediastinal lymph nodes in mice^38^. At day 21, we noted a severe reduction in splenic GC size, GC B cell proportions, and GC B cell counts relative to T_FH_ in B-Asns mice deprived of Asn (Fig. 5E-G, fig. S7F). Levels of anti-X31 IgG antibodies remained significantly lower in B-Asns mice fed an Asn-free diet, reproducing the findings at day 14 (Fig. 5H). Moreover, we found a robust numerical and proportional defect in mediastinal GC B cells in B-Asns mice on an Asn-free diet (Fig. 5I, fig. S7G). Additionally, there was a reduction in both the DZ to LZ and GC B to T_FH_ cell ratios, resembling the phenotype observed with ASNase treatment, but a non-significant change in mediastinal lymph node plasma cell numbers despite a decreasing trend (fig. S7H-J).

Asn metabolism therefore regulates GC-derived humoral immune responses during influenza infection, and *Asns* plays an indispensable role in this process when Asn availability is restricted.

### Disruption of Asn availability alters B cell metabolism

To understand how Asn withdrawal affects B cell metabolism, we performed stable isotope resolved LC-MS on B cells from B-WT or B-Asns mice, stimulated for 72h in Asn-replete media, before switching for 24h into media with or without Asn, and [U-^13^C]-glutamine, or ^15^N_1_-glutamine, labelled at either the amine or amide nitrogen position (Fig. 6A). Since ASNS transfers the amide nitrogen group of glutamine to that of asparagine (acting as an amidotransferase), positional labelling can provide information on ASNS activity^26^.

**Figure 6.**
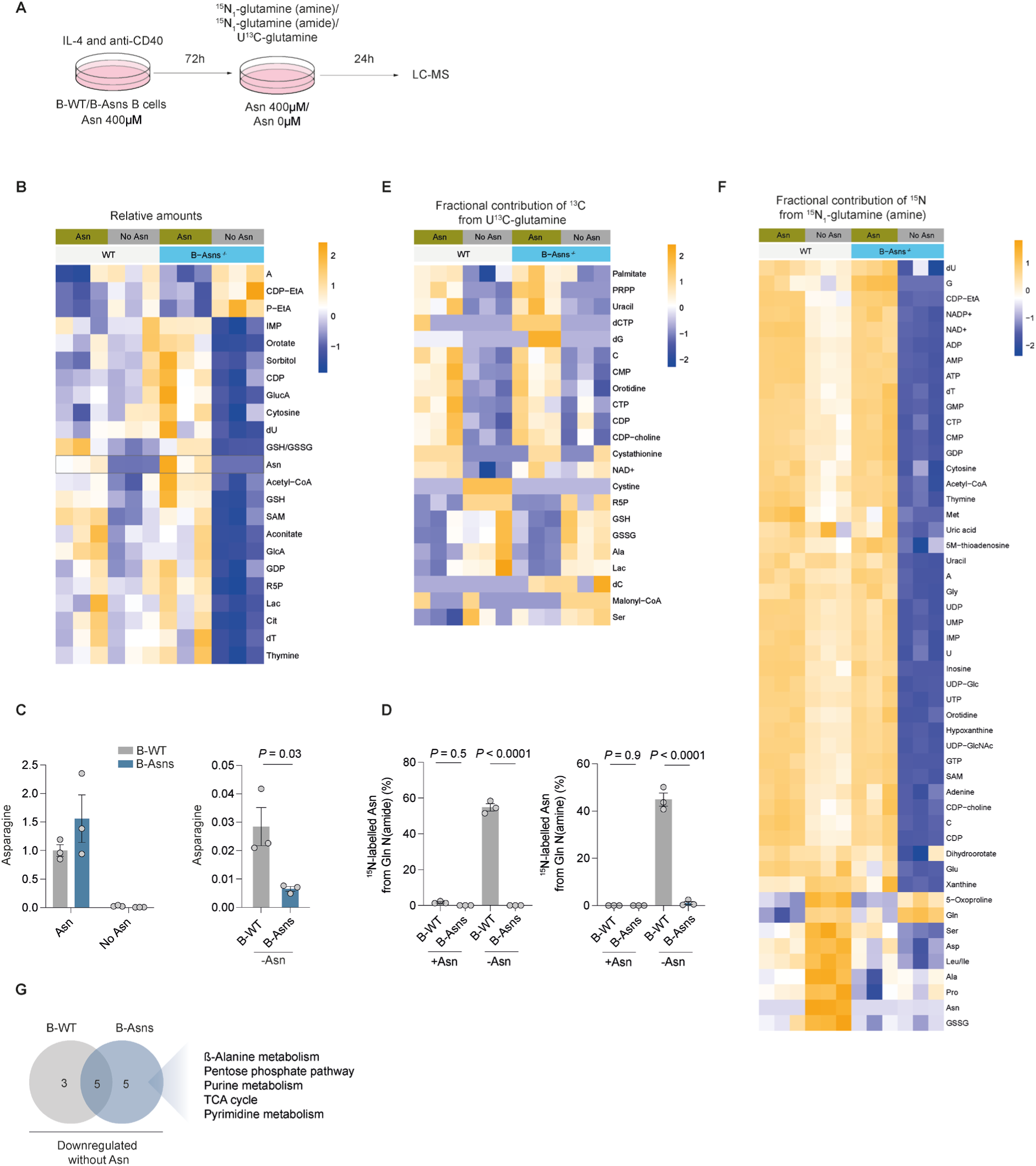
Disruption of asparagine availability alters B cell metabolism. **A.** Schematic of experimental design for stable isotope-resolved metabolomics. B-WT or B-Asns B cells were stimulated with IL-4 and agonistic anti-CD40 for 72 hours in the presence of Asn (400μM), then stimulated for an additional 24h with or without Asn (400μM), and either ^15^N_1_(amine)-glutamine, ^15^N_1_(amide)-glutamine, U^13^C-glutamine, or unlabelled glutamine, before analysis by LC-MS. **B.** Heatmap of total relative amounts of significantly different metabolites in B cells from B-WT or B-Asns mice as in **A**. Scale represents row Z-score. Significantly differentially abundant metabolites (P_adj_ < 0.05) are shown. Representative of two independent experiments. **C.** Abundance of intracellular Asn in B cells cultured as in **A**, and rescaled plot of Asn-deprived condition. Representative of two independent experiments. **D.** Fractional contribution of ^15^N to Asn pool, derived from ^15^N_1_-amino or amide Gln, following culture as described in **A**. **E.** Heatmap of fractional contribution of ^13^C to indicated metabolites, derived from U-^13^C-Gln following culture as described in **A**. Scale represents row Z-score. Significantly differentially labelled metabolites (P_adj_ < 0.05) are shown. **F.** Heatmap of fractional contribution of ^15^N to indicated metabolites, derived from ^15^N_1_-amine Gln following culture as described in **A**, with labelled compound added for final 24h of culture. Scale represents row Z-score. Significantly differentially labelled metabolites (P_adj_ < 0.05) are shown. **G.** Venn-diagram indicating number of significantly differentially-regulated pathways in B-WT and B-Asns B cells following Asn-withdrawal, based on relative metabolite abundance as in **B**. Listed pathways are those specifically downregulated in B-Asns B cells. Statistical significance was determined by two-way ANOVA with Šidák’s multiple testing correction (B,D-F), unpaired two-tailed t test (C), or hypergeometric test (G). Data are presented as the mean +/− SEM.

Intracellular Asn levels were greatly reduced in Asn-deprived conditions in both B-WT and B-Asns B cells, confirming our previous observation that exogenous uptake is dominant, but to significantly greater degree in those from B-Asns mice (Fig. 6B-C). B-WT cells did not engage in significant Asn synthesis when Asn was present, with negligible labelling of Asn with Gln-derived ^15^N from either the amino or amide position, despite ASNS expression (Fig 6D). Following Asn deprivation, around half of the intracellular Asn pool was ^15^N-labelled in B-WT B cells, reflecting de novo synthesis by ASNS. As expected, B-Asns B cells were incapable of Asn synthesis and no Asn was ^15^N labelled. There was minimal labelling of Asn from ^13^C glutamine, but around a quarter of Asp was labelled, with no difference between B-WT and B-Asns B cells, suggesting very little of the Gln-derived Asp pool was diverted to Asn synthesis (Fig. 6E, fig. S8A).

In B-WT cells, Asn deprivation had subtle effects on metabolite relative abundance, but significantly reduced fractional labelling from [U-^13^C]-glutamine and ^15^N_1_-glutamine. Despite small differences in intracellular Asn, B-Asns B cells exhibited profound metabolic dysfunction upon Asn withdrawal, with large reductions in metabolite relative amounts and glutamine-derived ^15^N incorporation (Fig. 6F).

We next performed pathway analysis to systemically describe these results. Comparing B-WT and B-Asns B cells, both deprived of Asn, there were a common set of significantly different pathways (Fig. 6G), including AAG, tRNA biosynthesis, glutathione synthesis, sphingolipid metabolism and glyoxylate. We noted that B-Asns B cells, when deprived of Asn, substantially downregulated metabolites in the TCA cycle, nucleotide biosynthesis. pentose phosphate pathway, and β-alanine metabolic pathways (Fig. 6G). Asn is therefore essential to regulate metabolic homeostasis in B cells.

### Mitochondrial function in B cells requires Asn

Given the abnormalities in TCA cycle metabolism we observed following Asn withdrawal, we next performed extracellular flux analysis to explore OxPhos and glycolysis in a dynamic setting. We found that Asn concentration was strongly associated with oxygen consumption rate (OCR) and extracellular acidification (ECAR), reflecting OxPhos and lactate production respectively (Fig. 7A-E, fig. S9A-B), and the deleterious effect of low Asn was more pronounced in B-Asns B cells (Fig 7C-D). In keeping with our metabolite profiling, this effect was only seen when Asn was limited. Surprisingly, the maximal respiration induced by the mitochondrial uncoupler carbonyl cyanide-p-trifluoromethoxyphenylhydrazone (FCCP) remained unchanged between B-WT and B-Asns B cells across all Asn concentrations, resulting in a significantly higher maximal-to-basal respiration ratio, known as the spare respiratory capacity (SRC), in B-Asns B cells (Fig. 7E, fig. S9B).

**Figure 7.**
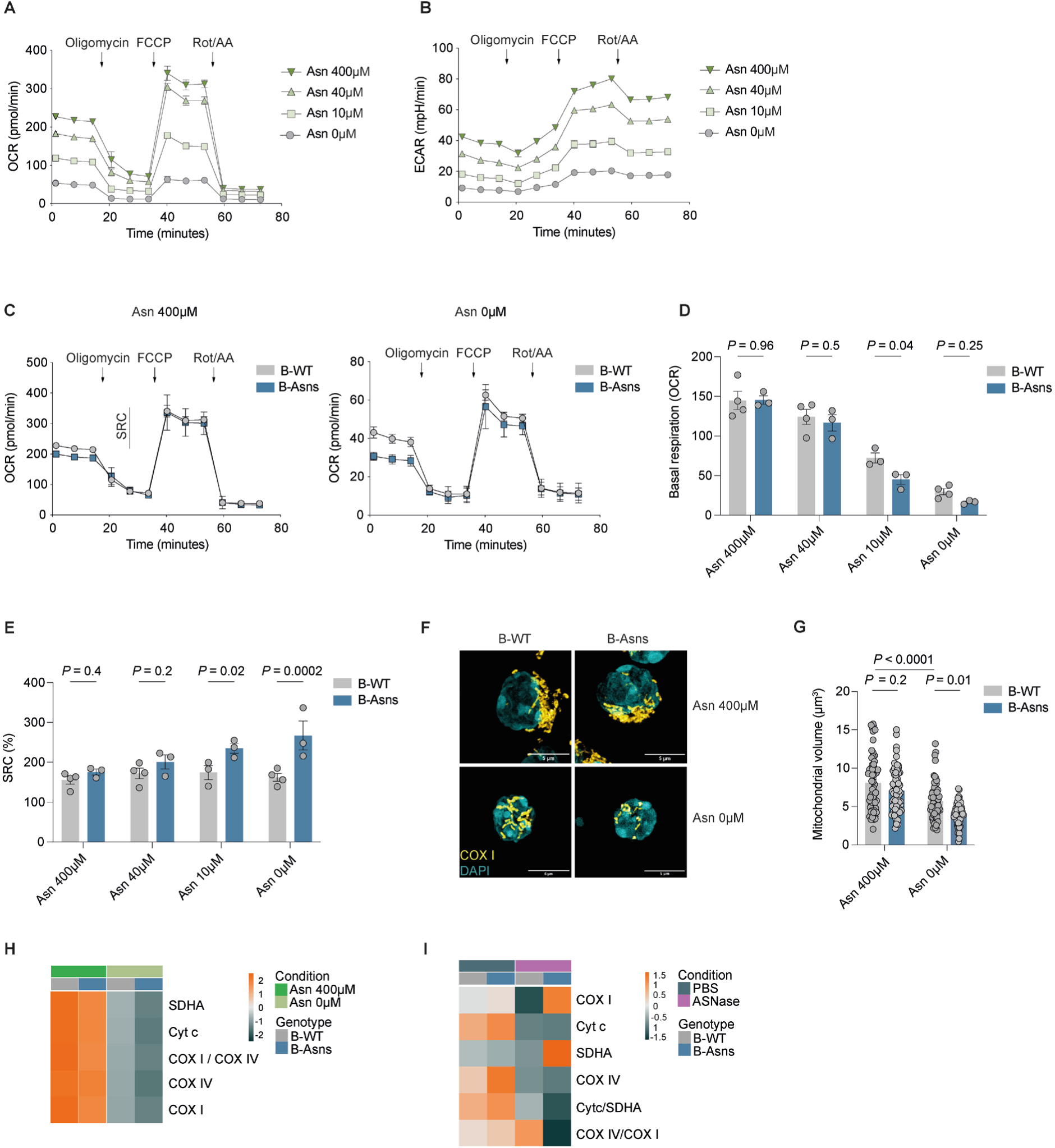
Mitochondrial function in B cells requires Asn. **A.** Seahorse extracellular flux analysis of B cells stimulated for 24h with IL-4 and agonistic anti-CD40 at the indicated concentrations of Asn. Oxygen consumption rate (OCR) was measured during the MitoStress test (n=3-4 mice). Data pooled from three independent experiments with 5-6 technical replicates averaged per mouse. FCCP, carbonyl cyanide-p-trifluoromethoxyphenylhydrazone; Rot/AA, rotenone/antimycin A **B.** Seahorse extracellular flux analysis of B cells as in **A** (n=3-4 mice). Extracellular acidification rate (ECAR) was measured during the MitoStress test. Data pooled from three independent experiments with 5-6 technical replicates averaged per mouse. **C.** Seahorse OCR quantification in B-WT (n=3-4 mice) and B-Asns (n=3 mice) B cells stimulated for 24h with IL-4 and agonistic anti-CD40 at the indicated concentrations of Asn. Spare respiratory capacity (SRC) is illustrated. Data pooled from three independent experiments. **D.** Quantification of basal OCR in B-WT and B-Asns B cells stimulated for 24h with IL-4 and agonistic anti-CD40 at the indicated concentrations of Asn. Each data point represents a mouse. Data pooled from three independent experiments. **E.** Quantification of SRC (%) based on quantifications in **fig. S9B** and Fig. 7D. **F.** 3D lattice SIM images of COX I and DAPI in B-WT and B-Asns B cells stimulated for 24h with IL-4 and agonistic anti-CD40 in the presence or absence of Asn (400μM). Scale bar 5µm. **G.** Quantification of mitochondrial volume as in **F**. Each data point represents a cell (n=57 B-WT/Asn400, n=66 B-Asns/Asn400, n=65 B-WT/Asn0, n=53 B-Asns/Asn0) pooled from n=3 mice B-WT or B-Asns mice. Representative of two independent experiments. **H.** Heatmap of row z-scores for the gMFI of indicated ETC proteins, measured by flow cytometry in B-WT or B-Asns B cells stimulated for 24h with IL-4 and agonistic anti-CD40 in the presence or absence of Asn (400μM) (mean of n = 2), Results are representative of two independent experiments with n = 3 mice per group in total. **I.** Heatmap of row z-scores for the gMFI of indicated ETC proteins and ratios, measured by flow cytometry in CD38^−^GL-7^+^ GC B cells from B-WT or B-Asns (mean of n = 3-4) mice treated with ASNase (standard regime). Data representative of two independent experiments. Statistical significance was determined by two-way ANOVA with Tukey’s multiple testing correction (D-E, G). Data are presented as the mean +/− SEM.

Supplementation with a cell-permeable analogue of the TCA cycle metabolite alpha-ketoglutarate (α-KG), dimethyl 2-oxoglutarate (DM-OG) has been reported to increase cell viability in T cells treated with L-asparaginase^39^. We therefore hypothesised that supplementation with DM-OG could potentially restore defective mitochondrial metabolism and improve B cell homeostasis in Asn-deficient conditions. However, DM-OG failed to rescue the defective B cell survival in B-WT and B-Asns B cells under Asn-free conditions (fig. S9C).

We next directly imaged mitochondria in B-WT and B-Asns B cells using 3D Lattice structured illumination microscopy (SIM) (Fig. 7F). Mitochondrial volume and count were not different when Asn was present, but were much reduced in the absence of Asn, and again this was seen to a greater extent in B-Asns B cells (Fig. 7G, fig. S9D).

The electron transport chain (ETC) is crucial for OxPhos activity. B cells undergo profound mitochondrial remodelling upon activation associated with ETC protein expression^2^. Using flow cytometry, we characterised the ETC protein abundances in activated B cells and found that Asn deprivation led to a substantial reduction in the expression of ETC proteins cytochrome c oxidase subunit I (COX I), cytochrome c oxidase subunit IV (COX IV), succinate dehydrogenase A (SDHA), and cytochrome c (Cyt c) in B-WT and to a greater extent in B-Asns B cells (Fig. 7H).

We next examined ETC proteins in *ex vivo* GC B cells from B-WT and B-Asns mice following ASNase. This again showed a decrease in levels of Cyt c and COX IV, but upregulation of SDHA and COX I, suggesting divergent contributions of Asn to mitochondrial regulation in GC B cells (Fig. 7I).

Taken together, these findings indicate that Asn availability maintains mitochondrial ETC homeostasis, and ASNS regulates mitochondrial respiration in low Asn conditions.

### Asn regulates nucleotide metabolism and the integrated stress response

Mitochondrial function is intimately linked to nucleotide metabolism, and in our GC B cell metabolomics experiment we found low levels of nucleotides, suggesting that their availability may be limiting, even in normal physiology (Fig. 1F)^40^. When examining the effect of Asn on B cell metabolite abundance, we found that nucleotide synthesis pathways were selectively impaired in B-Asns B cells following Asn deprivation (Fig. 6G). Detailed metabolite pathway analysis revealed significant downregulation of intermediate metabolites in both de novo purine and pyrimidine biosynthesis branches (fig. S10A). Orotate and uridine monophosphate (UMP), a common precursor of pyrimidine end-products cytosine and thymidine, were downregulated in B-Asns B cells.

The enzyme catalysing the oxidation of dihydroorotate to orotate is dihydroorotate dehydrogenase (DHODH), which is located in the mitochondria and forms a functional link between de novo pyrimidine biosynthesis and ETC activity^41,42^. We quantified DHODH levels and found it markedly reduced with Asn deprivation (Fig. 8A). Notably, B-Asns B cells had the lowest DHODH levels in the absence of Asn, consistent with their lower mitochondrial mass and significantly downregulated pyrimidine metabolites. Interestingly, 5-aminoimidazole-4-carboxamide ribonucleotide formyltransferase/IMP cyclohydrolase (ATIC), which is responsible for the synthesis of inosine monophosphate (IMP) in the de novo purine synthesis pathway was not altered at the protein level (Fig. 8A).

**Figure 8.**
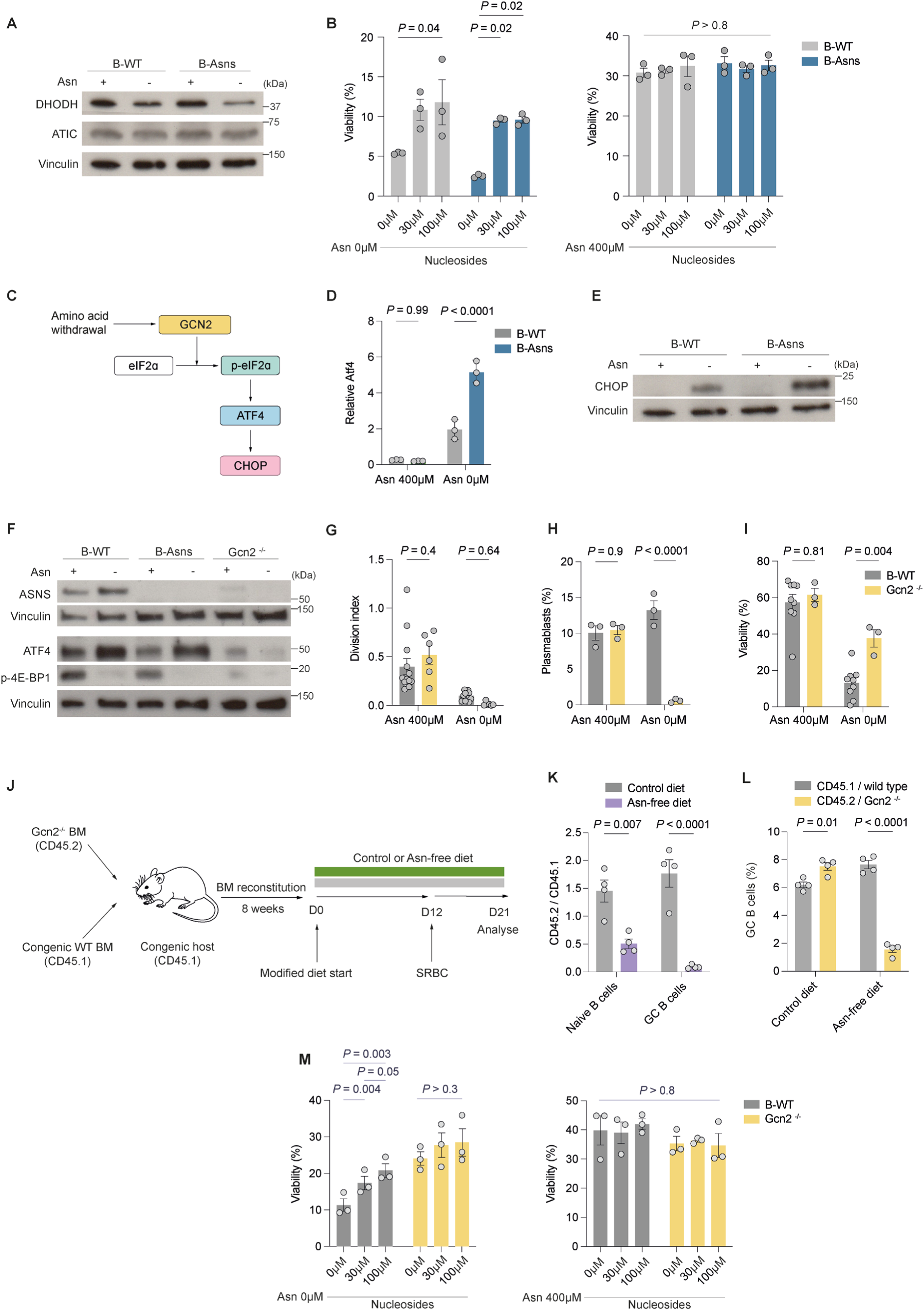
Asn regulates nucleotide metabolism and the integrated stress response. **A.** Representative immunoblot of DHODH and ATIC in B-WT or B-Asns B cells stimulated with IL-4 and anti-CD40 for 24h, in the presence or absence of Asn (400μM). Representative of two independent experiments. Vinculin used as loading control. **B.** Viability of B cells from B-Asns and B-WT mice (n=3 mice) stimulated for 72h with IL-4 and anti-CD40 in the presence or absence of Asn, supplemented with adenosine, thymidine, uridine, cytidine, and guanosine at 0μM, 30μM or 100μM each. Data pooled from two independent experiments. **C.** Schematic of the GCN2 branch of the integrated stress response. **D.** Relative expression of *Atf4* in B-Asns and B-WT B cells stimulated for 72h in the presence or absence of Asn (n=3 B-WT and n=3 B-Asns mice). Representative of two independent experiments. **E.** Representative immunoblot of CHOP in B-WT or B-Asns B cells stimulated with IL-4 and anti-CD40 for 24h, in the absence or presence of Asn (400μM). Representative of two independent experiments. Vinculin used as loading control. **F.** Representative immunoblot of ASNS, ATF4, phospho-4E-BP1 in B cells from B-Asns, *Gcn2*^−/−^, and B-WT mice, stimulated for 24h with anti-CD40 and IL-4 in the presence or absence of Asn (400μM). Representative of two independent experiments. Vinculin used as loading control. **G.** Division index of *Gcn2^−/−^* and wild type B cells stimulated with IL-4 and agonistic anti-CD40 for 72h, in the presence or absence of Asn (400μM). Each data point represents a mouse. Data pooled from >3 independent experiments. **H.** Representative flow cytometry quantification of plasmablast (IgD^−^ CD138^+^) differentiation of *Gcn2^−/−^* and wild type B cells following stimulation with LPS (10μg/ml) and IL-4 (50ng/ml) for 72h in the presence or absence of Asn (400μM) (n=3 *Gcn2^−/−^* mice and n=3 wild type mice). Data pooled from two independent experiments. **I.** Viability of *Gcn2^−/−^* and wild type B cells stimulated with IL-4 and agonistic anti-CD40 for 72h, in the absence or presence of Asn (400μM) (n=3 *Gcn2^−/−^* mice and n=9 wild type mice). Data pooled from >3 independent experiments. **J.** Schematic of bone marrow (BM) chimera/dietary modification experiment. Mixed BM chimeras were generated with wild type (CD45.1) and *Gcn2*^−/−^ (CD45.2) bone marrow. After an 8-week reconstitution period, mice were transferred to a control (Asn-replete) or Asn-deficient diet for 12 days, before immunisation with SRBC, similar to **fig. S5A**. Spleens were analysed 9 days after immunisation. **K.** CD45.2 *Gcn2*^−/−^ CD19^+^IgD^+^GL-7^−^ naïve B cell and CD38^−^BCL-6^+^ GC B cell abundance relative to CD45.1 wild type counterparts, fed control or Asn-free diets (n=4 mice). Data pooled from two independent experiments. **L.** CD38^−^BCL-6^+^ GC B cell frequencies in the CD45.2^+^*Gcn2*^−/−^ and CD45.1^+^ wild type B cell compartments under control or Asn-free diet (n=4 mice) settings. Data pooled from two independent experiments. **M.** Viability of B cells from B-WT or *Gcn2^−/−^* mice (n=3 mice) stimulated for 72h with IL-4 and agonistic anti-CD40 in the presence or absence of Asn, supplemented with adenosine, thymidine, uridine, cytidine, and guanosine at 0μM, 30μM or 100μM each. Data pooled from two independent experiments. Statistical significance was determined by repeated measure two-way ANOVA with Tukey’s multiple testing correction (B,D,K) or Šidák’s multiple testing correction (G-I,M), one-way ANOVA with Tukey’s multiple testing correction (L). Data are presented as the mean +/− SEM.

We next performed a rescue experiment by supplementing B-WT and B-Asns B cells with nucleosides including guanosine, adenosine, thymidine, cytidine, and uridine, which bypasses the activity of nucleotide biosynthetic enzymes, including DHODH. In conditions of Asn restriction, nucleoside supplementation substantially increased viability in both B-WT and B-Asns B cells following 72h culture (Fig. 8B), but was unable to restore normal proliferative capacity (fig. S10B). Importantly, nucleoside supplementation did not affect B cell viability or proliferation when Asn was unrestricted.

*Asns* is known to be upregulated following activation of the integrated stress response (ISR), which occurs as a consequence of a variety of cellular stressors, including amino acid withdrawal^43,44^. A key event in the amino acid response ISR is the phosphorylation of elongation initiation factor 2 alpha (eIF2ɑ) by the kinase general control non-repressible-2 (GCN2). Phosphorylated eIF2ɑ then controls transcription and translation of the ISR effector molecules activating transcription factor-4 (ATF4) and C/EBP homologous protein (CHOP)(Fig. 8C).

We examined the temporal dynamics of eIF2ɑ phosphorylation in B cells, and found that p-eIF2ɑ was increased at 6 hours following stimulation with IL-4 and agonistic anti-CD40 in the absence of Asn (fig. S10C). *Atf4* transcription was elevated with Asn deprivation, and this effect was significantly enhanced in B-Asns B cells (Fig. 8D). CHOP was highly expressed in B-Asns B cells deprived of Asn, and to a lesser extent in B-WT B cells, following *in vitro* activation with IL-4 and agonistic anti-CD40 (Fig. 8E). We observed a reduction in mTORC1 activation assessed by phosphorylation of 4E-BP1 and S6 when B-Asns B cells were stimulated without Asn (Fig. 8F, fig. S10D). This was accompanied by an increase in expression of the amino acid transporter component CD98 and SLC7A1 (fig. S10D). Interestingly, however, when we examined *ex vivo* GC B cells from mice treated with ASNase, whilst p-eIF2ɑ was elevated following ASNase in B-Asns mice, we also observed an unexpected increase in p-4E-BP1, p-S6, and the cell cycle regulator phosphorylated-Retinoblastoma protein (p-Rb) (fig. S10E).

To directly assess the importance of the GCN2-mediated ISR in Asn synthesis, we next examined *Gcn2*^−/−^ B cells. *Gcn2*^−/−^ B cells were unable to upregulate either ATF4 or ASNS following stimulation (Fig. 8F). Stimulation of *Gcn2*^−/−^ B cells resulted in a similar phenotype to B-Asns B cells, albeit with a paradoxical increase in viability in the absence of Asn (Fig 8G-I). To further investigate this finding, we then treated B cells with the GCN2 activator halofuginone^45^, and found that even when Asn was present, very low concentrations led to decreased survival (fig. S8F). This suggested that the extent of the ISR was carefully balanced in activated B cells, and excessive ISR activation was harmful. The GCN2 branch of the ISR is therefore essential for ASNS expression and cellular homeostasis in B cells.

We next examined the GC reaction in *Gcn2*^−/−^ mice, by immunisation with NP-CGG. At day 14, there was no difference in GC B cell numbers or anti-NP antibody levels compared with WT mice (fig. S10G-I). To understand whether GCN2 was required for the GC reaction in the setting of dietary Asn deprivation, we generated mixed bone marrow chimeric mice with CD45.1 wild type and CD45.2 *Gcn2^−/−^* cells (Fig. 8J). Following reconstitution, they were placed on a control or Asn-free diet, and then immunised with SRBC. We found profound out-competition of CD45.2^+^*Gcn2^−/−^* lymphocytes, and reduced GC B cells under dietary Asn deprivation (Fig. 8K-L, fig. S10J). Within the GC B cell compartment, there was disturbance of DZ/LZ partitioning, with a decrease in the DZ proportion (fig. S10K), similar to our previous findings in B-Asns GC B cells.

Finally, we examined whether nucleotide supplementation affected survival in *Gcn2^−/−^* B cells deprived of Asn. We found that unlike B-WT and B-Asns B cells, addition of nucleosides did not improve viability (Fig. 8M). This suggests that the effects seen with loss of GCN2 on B cell survival are distinct to those with ASNS alone, and nucleoside availability is not a major limiting factor in this setting.

These data show that *Asns* therefore essential to regulate B cell metabolic homeostasis, and if it cannot be obtained in sufficient amounts from the environment or synthesised by ASNS in response to activation of the ISR, there is impairment of nucleotide biosynthesis.

## Discussion

Here we show using integrated multiomics that Asn metabolism is upregulated in GC B cells and required for their homeostasis, and that deprivation of Asn either from the external environment or by loss of its synthesis strongly alters cellular metabolism, and in particular impairs OxPhos and nucleotide synthetic capacity.

We found that GC B cells exhibited high levels of protein synthesis *in vivo*, expressed the amino acid transporter ASCT2, and avidly took up Asn. Initial stimulation in the presence of Asn provided a protective effect against subsequent Asn withdrawal, as has also been observed in CD8^+^ T cells^46^, which we found was mediated in part by upregulation of ASNS, which was almost undetectable in naïve B cells. Although the effects of amino acid uptake on T cells is increasingly well described^11,47^, how this influences B cell biology is still poorly understood and requires further study.

Lack of Asn severely affected cellular metabolism in B cells, and in particular OxPhos. This was in striking contrast to reports in CD8^+^ T cells^13,15^, in which Asn deprivation leads to activation of OxPhos and increased nucleotide synthesis, associated with enhancement of proliferative capacity and enlarged mitochondrial mass, although others have noted reduction of cell division without Asn in this cell type, and also in cancer cells^12,14,48^. We found that Asn-deprivation led to a substantial drop in mitochondrial volume, and we observed that there was imbalance of expression of ETC proteins. It is unclear why B cells should be so much more vulnerable to the lack of Asn than CD8^+^ T cells, but may reflect more profound cellular remodelling following activation, or other differences in core metabolism. We did not observe a defect in T_FH_ cells with Asn depletion, despite their shared microenvironment with GC B cells, highlighting their divergent metabolism. However, we cannot fully exclude a more subtle functional defect.

An important consideration in all studies of metabolism *in vitro* is how to approximate the concentrations of metabolites found within tissue. Whilst this has received attention in the context of the tumour microenvironment, very little is known about normal tissues and especially within complex structures like GCs. GCs are hypoxic and poorly vascularised, but local concentrations of Asn remain unclear. We found a much higher concentration of Asn in eluted LNIF compared to serum, which was not the case for Gln. However, whether this applies in the complex microenvironment of the GC remains to be established. Advances in mass spectrometry imaging may allow more accurate quantification of metabolites in their spatial context in the future^49^.

Using mice lacking GCN2, we confirmed that activation of GCN2-mediated ISR is required for upregulation of *Asns*. Much like B-Asns mice, there was little effect of GCN2 deletion on the GC reaction with a normal diet. However, in a chimeric setting an Asn-free diet caused a severe defect in lymphocyte homeostasis, and a comparable defect in GC B cell numbers to similar B-Asns chimeras. This observation highlights the specific role for Asn metabolism during the GC reaction.

A key finding from our work is that Asn availability acts to gate nucleotide synthesis in B cells, a role previously demonstrated in cancer cell lines^50^. GC B cells have high rates of nucleic acid synthesis, in keeping with their active cell division, and our LC-MS data revealed they had low levels of nucleosides, suggestive of their consumption. Analysis of Asn synthesis from glutamine showed that even following activation, which was associated with upregulation of ASNS, the great majority of Asn was exogenously acquired. Nonetheless ASNS was essential for metabolic homeostasis and nucleotide synthesis when Asn is restricted, despite seemingly modest rates of Asn synthesis. This raises the interesting question of whether ASNS might have other, non-synthetic functions, as has been recently demonstrated for the enzyme phosphoglycerate dehydrogenase (PHGDH), whose canonical function is to synthesise serine^51^. Using single cell RNA sequencing, we identified dysregulation of cell cycle gene expression pathways when Asn was deficient, and confirmed that whilst cyclin B1 was upregulated, mitosis was defective in B-Asns DZ GC B cells. Interestingly, ASNS has been found to be recruited to the mitotic spindle in dividing human cells under Asn deprivation although its function is unknown^52^. The precise role of ASNS in cell cycle progression and division therefore warrants further investigation.

The relationship between amino acid availability and nucleotide synthesis has been previously defined through mTORC1 signalling, acting via phosphorylation of the enzyme carbamoyl-phosphate synthetase 2, aspartate transcarbamoylase, and dihydroorotase (CAD), or the tetrahydrofolate cycle. It is therefore possible that in GC B cells, Asn availability tunes these nucleoside synthetic pathways, either through mTORC1 or other mechanisms. An important node in B cell nucleoside metabolism downregulated with Asn deprivation involved the enzyme which oxidises dihydroorotate, DHODH. However, the effect of Asn deprivation on mTORC1 signalling in B cells appears to vary according to setting. *In vitro* we found reduction in phosphorylation of mTORC1 target proteins. However, *ex vivo* we were surprised to find an increase following ASNase treatment in B-Asns GC B cells, accompanied by elevated phospho-Rb levels. The cause for this divergence is unclear, but may be related to differing environmental milleu, stimulation, or relative level of Asn deficiency. Whether this mTORC1 upregulation is maladaptive remains to be established.

ASNase has been a cornerstone of the treatment of leukaemia for decades, but given the results of our work, ASNS may also be a novel therapeutic target in non-malignant, autoimmune disease mediated by the GC reaction, and is deserving of future study in this context. Another aspect of ASNase is its role in promoting bacterial infection. It is interesting to speculate that its capacity to reduce environmental Asn is also effective against the protective humoral response, which is partially protected against by the expression of ASNS.

## Supplementary material

**Supplementary Figure 1.**
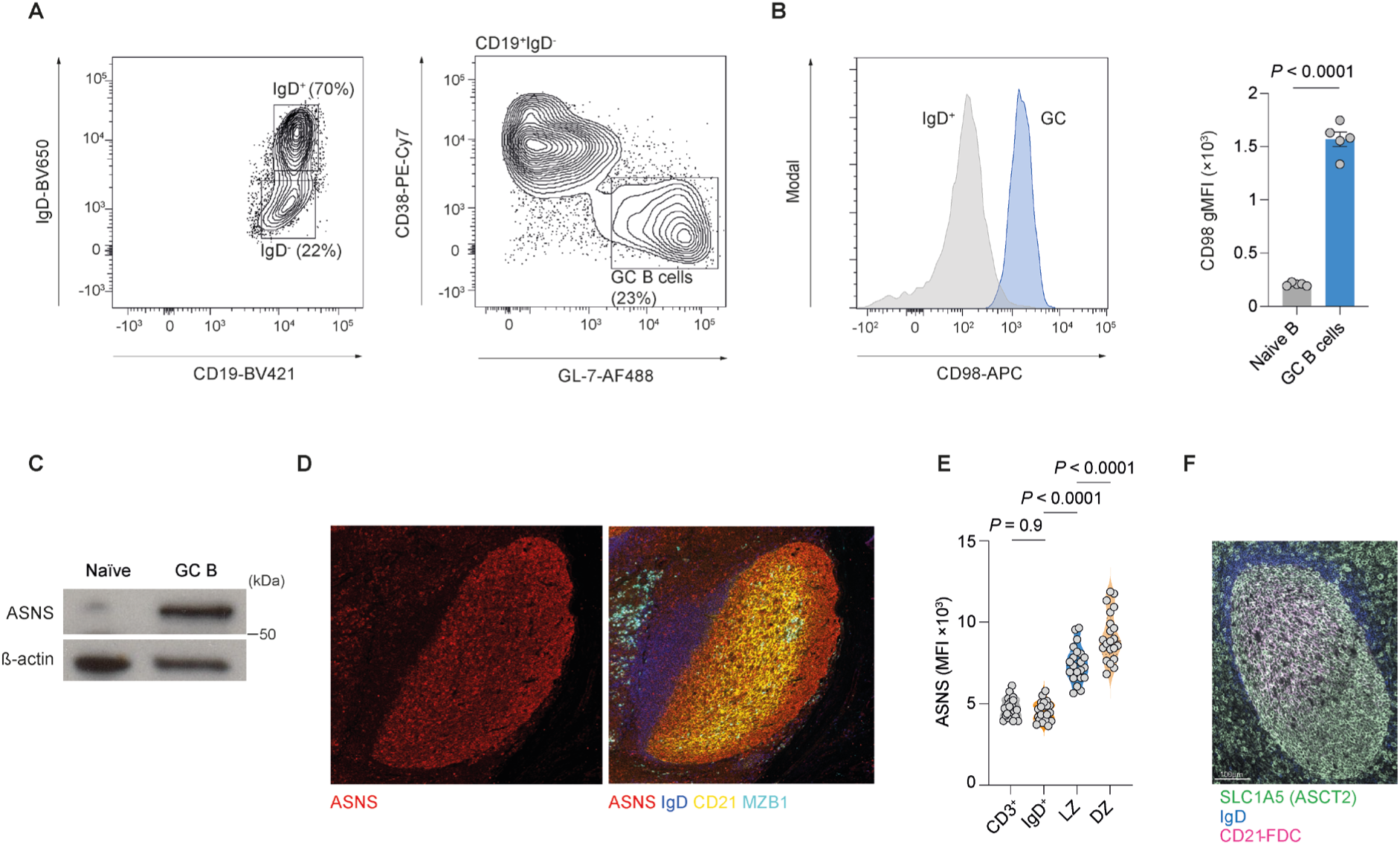
**A.** Gating strategy for GC B cells (CD19^+^IgD^lo^CD38^−^GL-7^+^) and naïve IgD^+^ B cells (CD19^+^IgD^+^). **B.** Representative flow cytometry histogram and quantification of CD98 geometric mean fluorescence intensity (gMFI) in IgD^+^ naïve and GC B cells (n=5 mice), as in **A**. Data representative of two independent experiments. **C.** Immunoblot of ASNS in magnetic bead sorted GC B cells and naïve B cells, from wild type mice immunised with SRBC at day 0 and day 4, with analysis at day 12. Representative of two independent experiments. Beta-actin is used as a loading control. **D.** CellDIVE immunofluorescence images of a GC from a tonsil highlighting ASNS in CD21^+^ LZ, CD21^−^ DZ, IgD^+^ naïve, and MZB1^+^ plasma cell niches. **E.** Quantification of mean fluorescence intensity (MFI) of ASNS in areas shown in **D,** including CD3^+^ T cells. Each dot represents a randomly-selected GC with a representative DZ, LZ and surrounding IgD^+^ naïve B cell follicle (n=21 GCs in total, pooled from three tonsils). Representative of two independent experiments. **F.** CellDIVE immunofluorescence images of a GC from a tonsil highlighting SLC1A5 (ASCT2) expression in CD21^+^ LZ, CD21^−^ DZ, and surrounding IgD^+^ naïve B cell follicle. Scale bar 100µm. Statistical significance was determined by unpaired two-tailed t test (B) or one-way ANOVA with Tukey’s multiple testing correction (E). Data are presented as the mean +/− SEM.

**Supplementary Figure 2.**
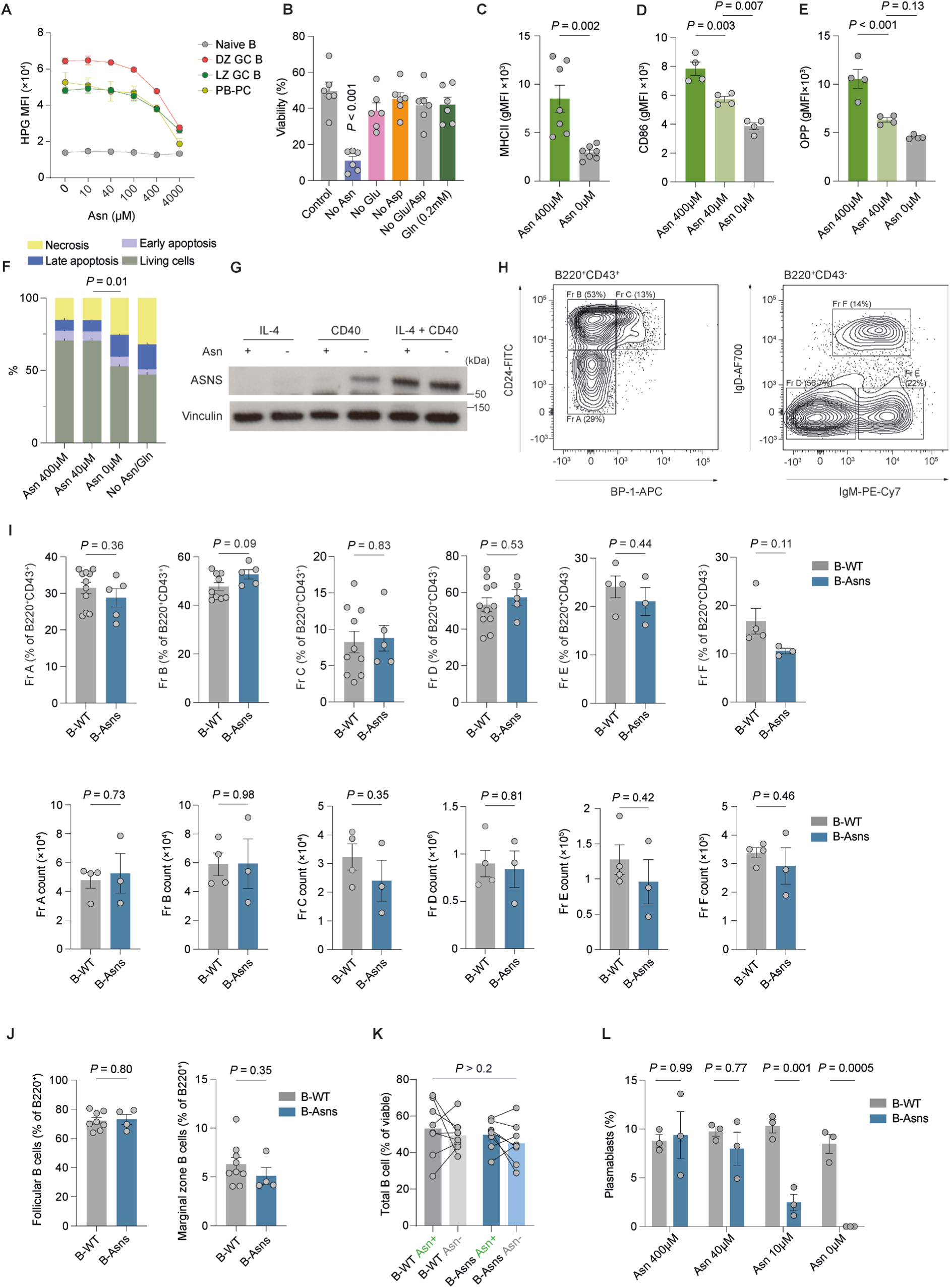
**A.** Quantification of HPG (400μM) uptake at increasing concentrations of Asn (0-4000μM) in splenic IgD^+^GL-7^−^ naïve B, IgD^−^CD138^+^ plasmablast/plasma cells (PB-PC), CD38^−^GL-7^+^CXCR4^lo^CD86^hi^ LZ and CD38^−^GL-7^+^CXCR4^hi^CD86^lo^ DZ GC B cells harvested from n=4 mice at day 8 following SRBC immunisation. Background signal subtraction was performed based on 4°C and no-HPG controls. Representative of two independent experiments. **B.** Viability of B cells stimulated for 72h with IL-4 and anti-CD40, in the presence or absence of the indicated amino acids (n=6 mice). Data pooled from three independent experiments. **C.** Major histocompatibility complex II (MHCII) gMFI in B cells stimulated for 24h with IL-4 and anti-CD40 with the indicated concentrations of Asn (n=7 mice). Data pooled from three independent experiments. **D.** CD86 gMFI in B cells stimulated for 24h with IL-4 and anti-CD40 with the indicated concentrations of Asn (n=4 mice). Representative of two independent experiments. **E.** OPP incorporation determined by flow cytometry in B cells stimulated for 24h with IL-4 and anti-CD40 with the indicated concentrations of Asn (n=4 mice). Representative of two independent experiments. **F.** Quantification of stages of apoptosis in B cells stimulated with IL-4 and anti-CD40 at 18h, defined by annexin V labelling and propidium iodide (PI) staining, in the indicated concentrations of Asn of Asn or Asn and Gln (n=3 mice). Early apoptotic: Annexin V^+^ PI^−^, late apoptotic: Annexin V^+^ PI^+^, necrotic: Annexin V^−^ PI^+^, live cells: Annexin V^−^ PI^−^. Representative of two independent experiments. **G.** Representative immunoblot of ASNS in B cells stimulated for 48h with IL-4, anti-CD40, or IL-4 and anti-CD40, in the presence/absence Asn. Representative of two independent experiments. **H.** Representative gating strategy for indicated Hardy stages of B cell development in bone marrow. **I.** Frequency and absolute count quantifications of indicated Hardy stages of B cell development in bone marrow as in **H**. Each data point represents a mouse. Data pooled from two independent experiments. **J.** Marginal zone and follicular B cell quantification in spleen. Each data point represents a mouse. Data pooled from two independent experiments. **K.** Quantification of CD19^+^ total B cell proportions by flow cytometry in lymph node slices (300μm thick sections) cultured for 20h in the presence or absence of Asn (400μM) obtained from inguinal lymph nodes of B-WT and B-Asns mice immunised with NP-CGG (day 14). Each dot represents a single lymph node. Data pooled from two independent experiments. **L.** Quantification of IgD^−^ CD138^+^ Plasmablast proportions in B cells from B-WT and B-Asns mice stimulated with LPS and IL-4 for 72h in the absence of presence of the indicated concentrations of Asn. Representative of three independent experiments. Statistical significance was determined by one-way ANOVA with Tukey’s (D-E) or Dunnett’s (B) multiple testing correction, unpaired two-tailed t test (C,I-J), paired two-tailed t test (K), or two-way ANOVA with Šidák’s multiple testing correction (L) or Tukey’s multiple testing correction (F). Data are presented as the mean +/− SEM.

**Supplementary Figure 3.**
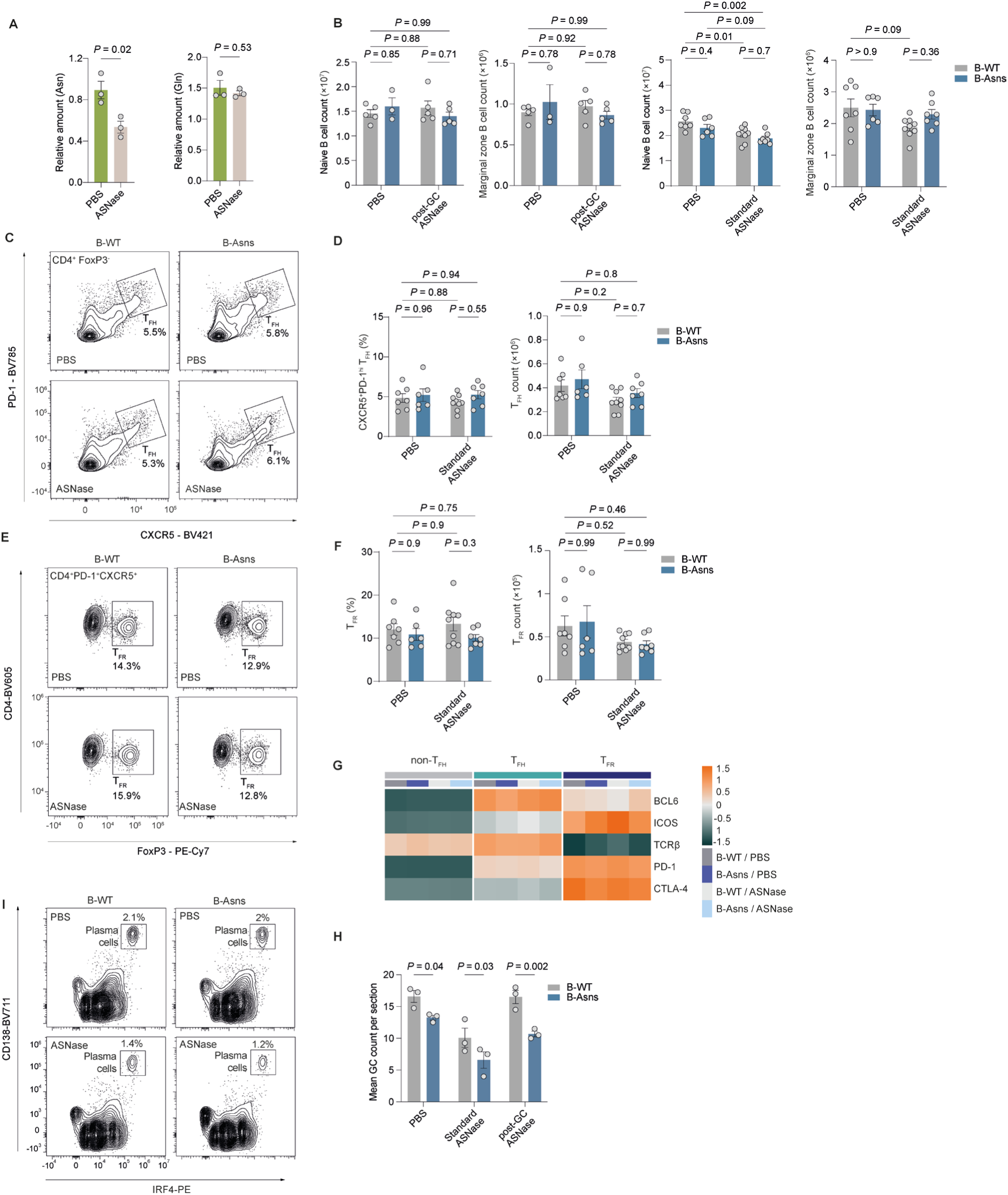
**A.** Serum Asn and Gln LC-MS quantification. WT mice were intraperitoneally injected with ASNase (500U/kg: ≈10U/mouse) or vehicle PBS, and 48h later terminally bled and sera were collected for LC-MS quantification of Asn and Gln (n=3 pools of 1-2 wild type mice, with control samples as in Fig. 2A). Representative of two independent experiments. **B.** Absolute count quantification of splenic naïve B cell (IgD^+^ GL-7^−^) and MZB (CD19^+^IgD^lo^ CD38^hi^, gating strategy independently validated and yielded >95% CD21/35^+^ CD23^−^ IgM^hi^ bona-fide MZB) from B-WT or B-Asns mice treated with PBS, standard or post-GC ASNase. Each point represents a single mouse. Data pooled from >2 independent experiments. **C.** Representative gating strategy for CD19^−^ CD4^+^ Foxp3^−^ CXCR5^hi^ PD-1^+^ T_FH_ cells **D.** Quantification of T_FH_ numbers and frequency (% of CD19^−^ CD4^+^ Foxp3^−^) at day 9 post SRBC immunisation following standard ASNase or PBS treatment regime. Each point represents a single mouse. Data pooled from three experiments. **E.** Representative gating strategy for CD19^−^CD4^+^CXCR5^+^PD-1^+^Foxp3^+^ T_FR_ cells **F.** Quantification of T_FR_ ratio (% of CD19^−^CD4^+^CXCR5^+^PD-1^+^) and counts at day 9 post SRBC immunisation following standard ASNase or PBS treatment regime. Each point represents a single mouse. Data pooled from three experiments. **G.** Heatmap of row z-scores for the gMFI values of indicated functional TFH-TFR proteins (BCL-6 and CTLA4 measured intracellularly) in non-TFH (CD19^−^ CD4^+^ PD-1^−^ CXCR5^−^), TFH (CD19^−^ CD4^+^ FoxP3^−^ CXCR5^+^ PD-1^+^), TFR (CD19^−^ CD4^+^ FoxP3^+^ CXCR5^+^ PD-1^+^), measured by flow cytometry at day 9 SRBC immunisation following post-GC ASNase regime (mean of n = 4-5 mice each condition). Data are pooled from two independent experiments. **H.** Quantification of GC numbers per spleen sections (average of at least two non-serially cut sections). Each point represents a mouse. Data pooled from two independent experiments. **I.** Representative gating strategy for plasma cells as IgD^−^ IRF4^+^ CD138^+^. Statistical significance was determined by unpaired two-tailed t test (A), two-way ANOVA with Tukey’s multiple testing correction (B,D,F) or Šidák’s multiple testing correction (H). Data are presented as the mean +/− SEM.

**Supplementary Figure 4.**
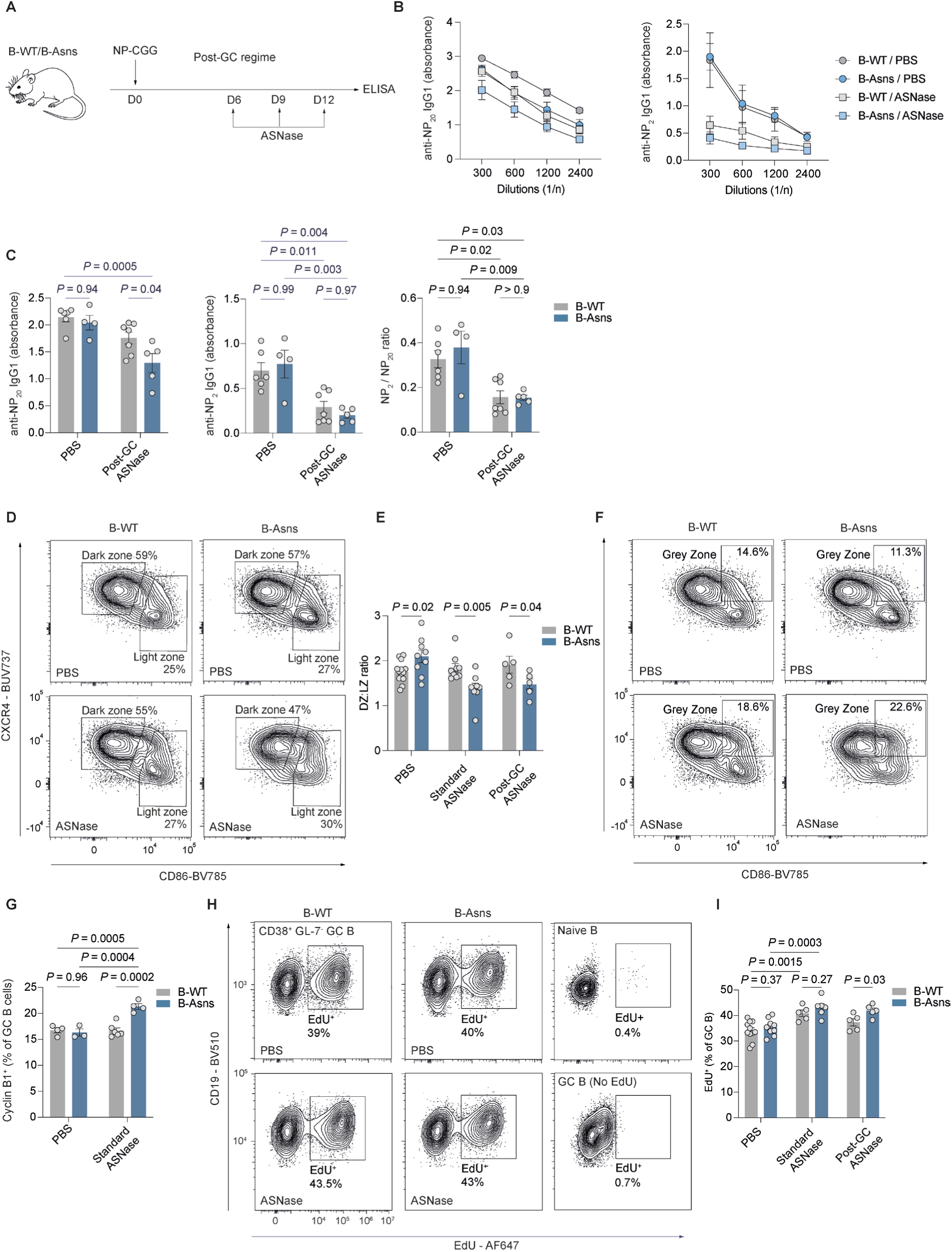
**A.** Schematic of in vivo ASNase administration regime for affinity maturation detection. B-WT or B-Asns mice were subcutaneously injected with 50μg NP-CGG precipitated in Alum hydrogel (1:1) subcutaneously on both flanks and given three ASNase (500U/kg: ≈10U/mouse) or PBS injections at day 6, 9, and 12 followed by serum collection at day 14 for ELISA. **B.** Dilution curves of IgG1 anti-NP antibodies (NP_>20_-BSA and NP_2_-BSA respectively) at day 14, n=4-7 mice each condition. Data pooled from three independent experiments. **C.** ELISA quantification of IgG1 anti-NP antibodies (NP_>20_-BSA and NP_2_-BSA at 1:300 dilution) and NP_2_:NP_20_ ratio at day 14, n=4-7 mice each condition. Data pooled from two independent experiments. **D.** Representative flow cytometry plots of light zone (CD86^hi^CXCR4^lo^) and dark zone (CD86^lo^CXCR4^hi^) GC B cells **E.** Quantification of DZ:LZ ratio. Data pooled from >3 independent experiments. **F.** Representative flow cytometry plots of grey zone (CD86^hi^CXCR4^hi^) GC B cells **G.** Quantification of Cyclin B1+ GC B cells following standard ASNase regime or PBS control. Data representative of two independent experiments. **H.** B-WT or B-Asns mice were treated with standard or post-GC ASNase regimes as described in Fig. 3A. On day 9 post-SRBC immunisation, mice were pulsed with 1mg EdU intraperitoneally and two hours later EdU incorporation in splenic GC B cells were analysed by flow cytometry. Representative gating strategy EdU^+^ GC B cell fraction with negative controls. **I.** Quantification of **H.** Data pooled from four independent experiments. Statistical significance was determined by two-way ANOVA with Šidák’s multiple testing correction (C,G) or Tukey’s multiple testing correction (E,I). Data are presented as the mean +/− SEM.

**Supplementary Figure 5.**
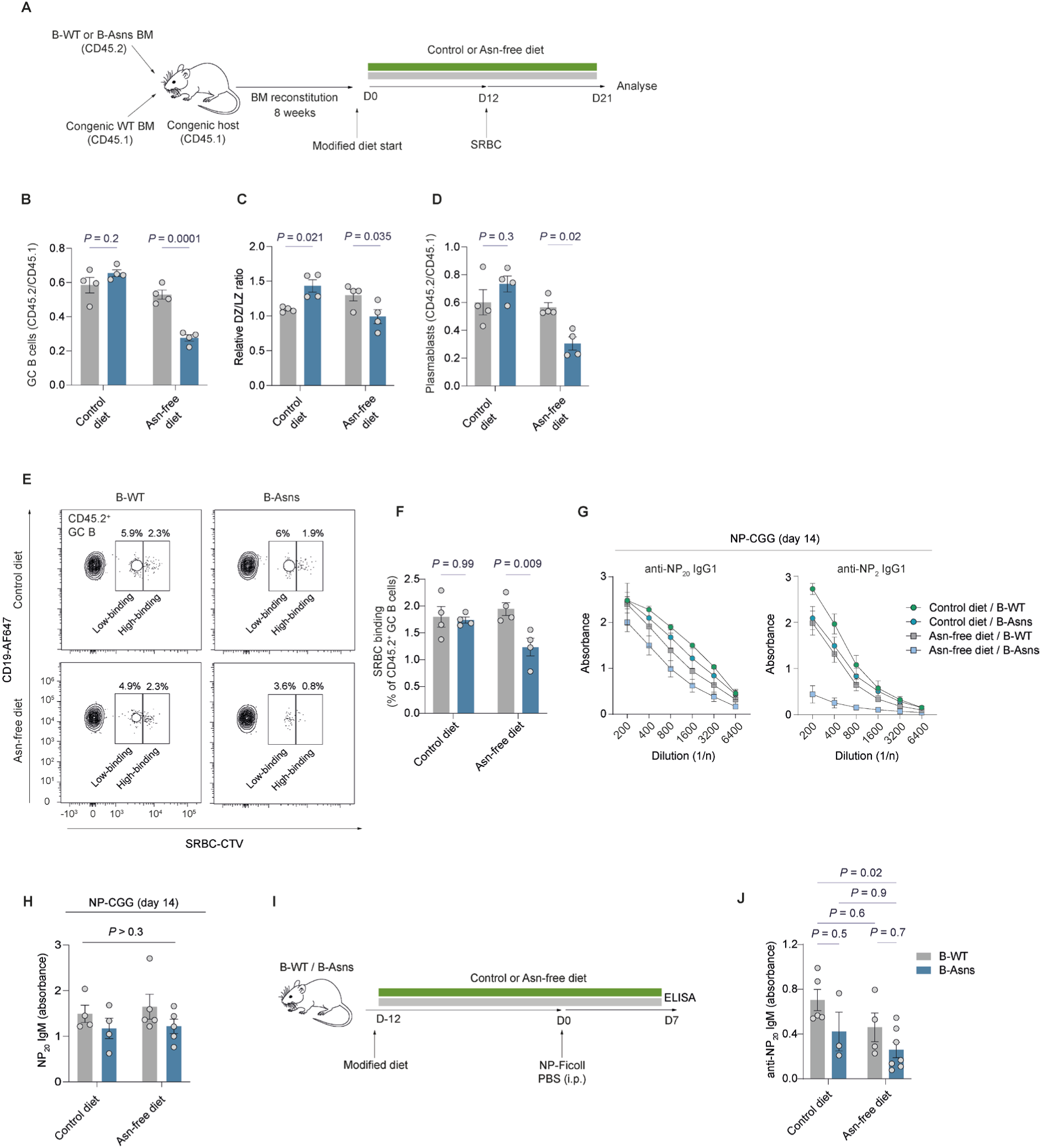
**A.** Schematic of bone marrow (BM) chimera/dietary modification experiment. Mixed BM chimeras were generated with B-WT (CD45.2) or B-Asns (CD45.2) and CD45.1 wild type BM, with n=4 congenic CD45.1 recipient mice for each condition. After an 8 week reconstitution period, mice were transferred to a control (Asn-replete) or Asn-deficient diet for 12 days, before immunisation with SRBC. Spleens were analysed 9 days after immunisation. **B.** Ratio of splenic GC B cells in CD45.2^+^ B cells normalised to CD45.1^+^ WT counterparts from bone marrow chimeric mice as described in **A**. Each data point represents a single mouse. Data pooled from two independent experiments. **C.** Ratio of DZ:LZ in CD45.2^+^ GC B cells normalised to CD45.1^+^ WT counterparts from bone marrow chimeric mice as described in **A**. Data pooled from two independent experiments. **D.** Ratio of splenic plasma cells (CD138^+^IRF4^+^) in CD45.2^+^ cells normalised to CD45.1^+^ WT counterparts from bone marrow chimeric mice as described in **A**. Data pooled from two independent experiments. **E.** Representative high and low binding fractions of CD45.2 GC B cells from BM chimeric recipient mice to SRBCs labelled with Celltrace Violet (CTV). **F.** Ratio of splenic CD45.2^+^ SRBC-binding (high-binding as described in **E**) GC B cells from bone marrow chimeric mice as described in **A.** Data pooled from two independent experiments. **G.** Dilution curves of IgG1 anti-NP antibodies (NP_>20_-BSA and NP_2_-BSA respectively) at day 14, n=4 mice each condition. Data pooled from two independent experiments. **H.** ELISA quantification of anti-NP_20_ IgM antibodies at day 14 post-NP-CGG immunisation. Each point represents a single mouse. Data pooled from two independent experiments. **I.** Schematic of NP-Ficoll/dietary modification experiment. B-WT or B-Asns mice were assigned to either an Asn-free or control diet (Asn-replete) for 12 days, before immunisation with 50μg NP-Ficoll in PBS (1:1) intraperitoneally. Mice were bled on day 7 for ELISA detection. **J.** ELISA quantification of anti-NP_20_ IgM antibodies at day 7 post-NP-Ficoll immunisation. Each point represents a single mouse. Data pooled from two independent experiments. Statistical significance was determined by two-way ANOVA with Šidák’s multiple testing correction. Data are presented as the mean +/− SEM.

**Supplementary Figure 6.**
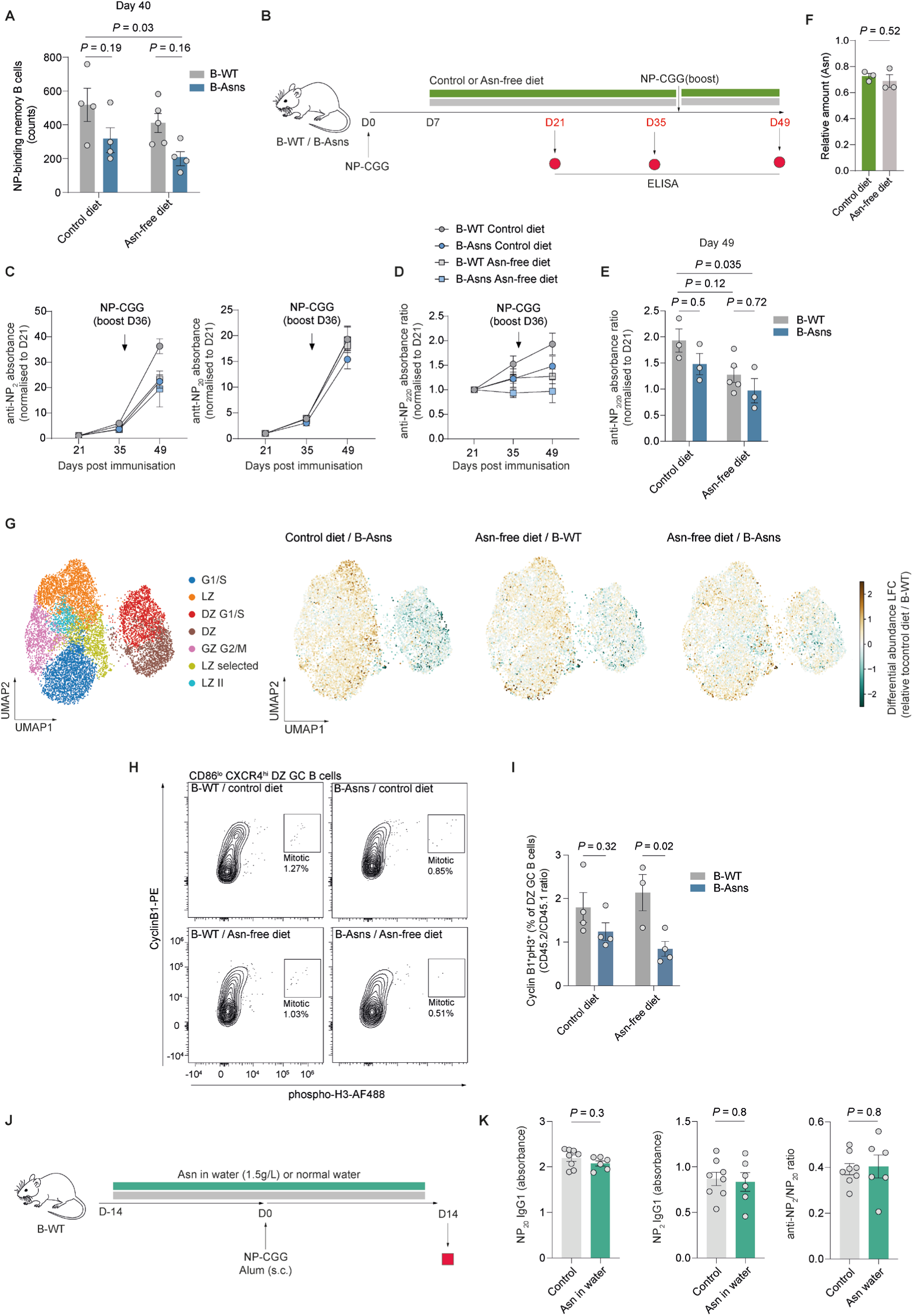
**A.** Quantification of NP-binding memory B cell counts in spleen at day 40 post-NP-CGG immunisation as in Fig. 4A. Data pooled from two independent experiments **B.** Schematic of NP-CGG/dietary modification experiment. B-WT or B-Asns mice immunisation with 50μg NP-CGG precipitated in Alum hydrogel (1:1) subcutaneously on both flanks, one week later, they were assigned to either an Asn-free or control diet (post-GC diet treatment) and maintained on the assigned diet for the rest of the experiment. Sera were collected on D21 and D35 before boost immunisation was given with 50μg NP-CGG precipitated in Alum hydrogel (1:1) subcutaneously on only right flank at D36. Two weeks later, mice were bled at day 49 timepoint and experiment was terminated. **C.** ELISA quantification of anti-NP_2_ IgG1, anti-NP_20_ IgG1 antibodies and NP_2_/NP_20_ ratio at day 21, 35 and 49 after primary NP-CGG immunisation (normalised to day 21). Each point represents a serum sample from single mouse. Data pooled from two independent experiments **D.** Anti-NP_2_/NP_20_ ratio quantification at day 21, 35 and 49 after primary NP-CGG immunisation (normalised to day 21). Each point represents a serum sample from single mouse. Data pooled from two independent experiments **E.** Comparison of normalised NP_2_/NP_20_ ratio between conditions at day 49 based on **D**. **F.** Serum samples collected as in **E** (42 days after diet start) and analysed by LC-MS (n=3 pools of 2-3 mice) for relative amount of Asn and Gln. Representative of two independent experiments. **G.** MrVI differential abundance log fold change (LFC) comparison across conditions relative to control diet / B-WT **H.** Representative flow gating of Cyclin B1+ phospho-Histone3^+^ mitotic cells in CD45.2 CXCR4^hi^ CD86^lo^ (DZ) GC B cells from BM chimeric recipient mice as in **fig S5A**. **I.** Ratio of splenic CD45.2^+^ DZ mitotic GC B cells (Cyclin B1+ phospho-Histone3^+^, as depicted in **H**) normalised to CD45.1^+^ WT counterparts from bone marrow chimeric mice as described in **fig S5A**. Data pooled from two independent experiments. **J.** Schematic of NP-CGG/Asn supplementation in water experiment. B-WT mice were administered normal or Asn supplemented-water (1.5g/L) for 14 days before subcutaneous immunisation with NP-CGG. Sera were collected at day 14 post-immunisation for the detection of NP-specific antibodies by ELISA. **K.** ELISA quantification of anti-NP_2_ and anti-NP_20_ IgG1 antibodies along with the NP_2_/NP_20_ ratio (affinity maturation index) at day 14 post-NP-CGG immunisation. Each point represents a single mouse. Data pooled from two independent experiments. Statistical significance was determined by two-way ANOVA with Šidák’s multiple testing correction (A, E, I) and unpaired two-tailed t test (F, K). Data are presented as the mean +/− SEM.

**Supplementary Figure 7.**
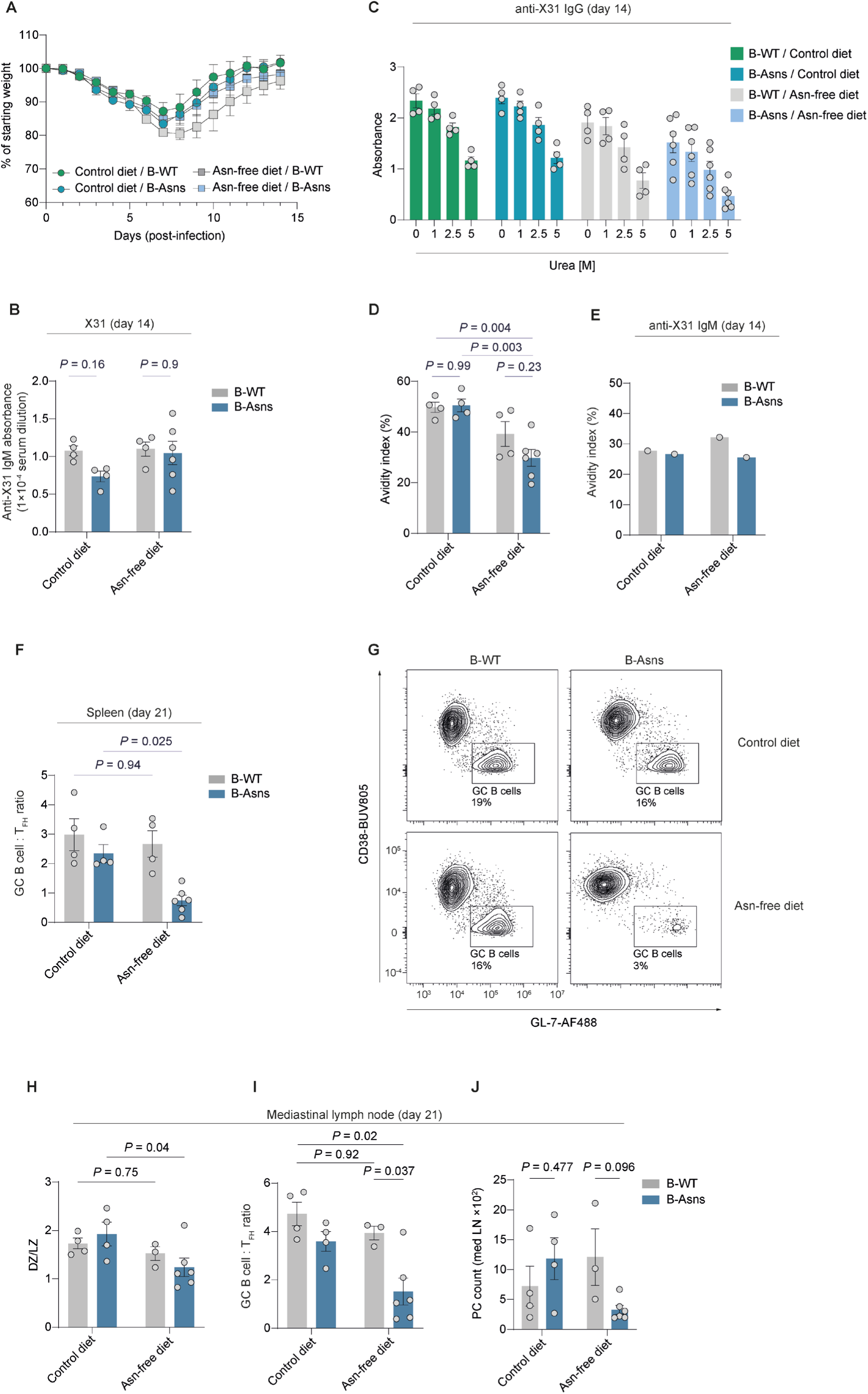
**A.** Quantification of weight loss as % of initial weight over 14 days. Control diet/ B-WT (n = 4), control diet/B-Asns (n = 4), Asn-free diet/B-WT (n = 4), Asn-free diet/B-Asns (n=6). Data pooled from two independent experiments. **B.** Quantification of anti-X31 IgM at day 14 following influenza infection, as depicted in Fig. 5A **C.** ELISA quantification at day 14 for anti-X31 IgG absorbance at 1/5×10^4^ dilution following increasing concentration of urea treatment (at 0, 1, 2.5 or 5M). Each point represents a single mouse. Data pooled from two independent experiments. **D.** Comparison of avidity (functional affinity) of X31-specific IgG at day 14 which was calculated as following: Absorbance [5M urea] / Absorbance [0M urea] × 100 also indicated in **C**. Data pooled from two independent experiments. **E.** Avidity index quantification for anti-X31 IgM based on 5M Urea treatment calculated as in **D**. Each point represents a pooled sera from n=4-6 mice per condition. Data representative of two independent experiments. **F.** Quantification of splenic GC B:T_FH_ ratio at day 21 post-influenza infection (n=4-6 mice per condition). Data pooled from two independent experiments. **G.** Representative flow cytometry plots of GC B cells in mediastinal lymph node at day 21. **H.** Quantification of DZ/LZ ratio in CD38^−^ GL-7^+^ GC B cells in mediastinal lymph node at day 21 post-influenza infection. Each point represents a single mouse (n=3-6 mice each condition). Data pooled from two independent experiments. **I.** Quantification of GC B:T_FH_ ratio in mediastinal lymph node at day 21 post-influenza infection (n=3-6 mice each condition). Data pooled from two independent experiments. **J.** Quantification of IgD^−^CD138^+^ plasma cell count in mediastinal lymph nodes at day 21. n=3-6 mice each condition. Data pooled from two independent experiments Statistical significance was determined by two-way ANOVA with Šidák’s (B,H-J) or Tukey’s (D,F) multiple testing correction. Data are presented as the mean +/− SEM.

**Supplementary Figure 8.**
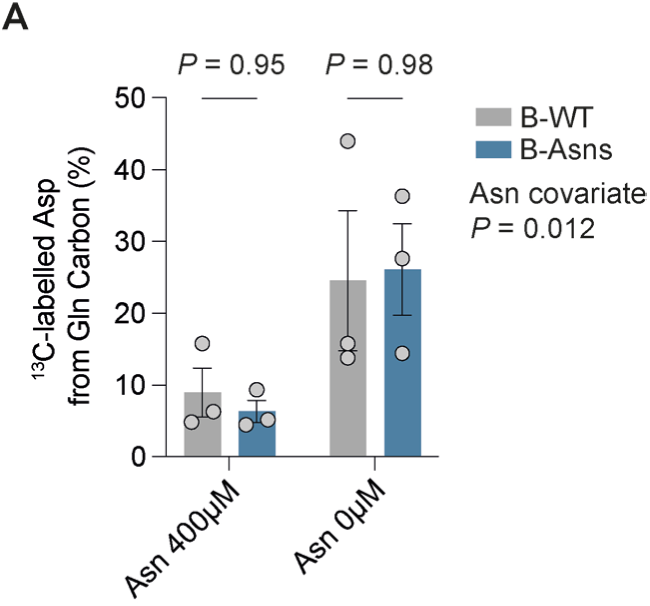
**A.** Percentage of aspartate labelled with ^13^C, indicating derivation from U^13^C-glutamine, as in Fig. 6E. Statistical significance was determined by two-way ANOVA with Šidák’s multiple testing correction, with Asn concentration as a co-variate. Data are presented as the mean +/− SEM.

**Supplementary Figure 9.**
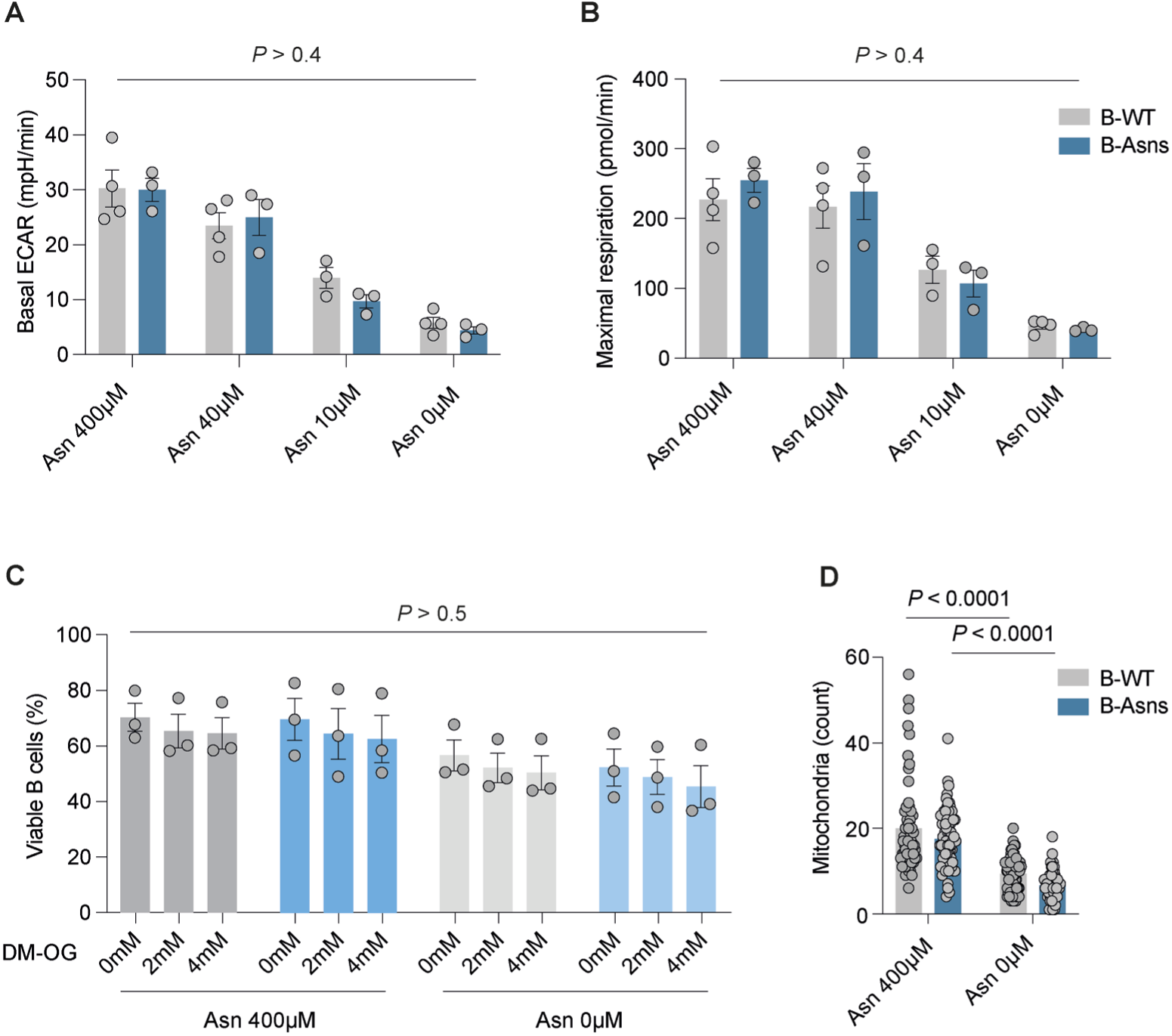
**A.** Quantification of basal ECAR as in Fig. 7B. **B**. Quantification of maximal respiration as in Fig. 7D. **C.** Viability of B cells from B-WT and B-Asns stimulated with anti-CD40 and IL-4 for 24h in the presence or absence of Asn (400μM), supplemented with vehicle (DMSO) or DM-OG at the indicated concentrations (0mM, 2mM or 4mM). Data pooled from two independent experiments (n=3 B-WT, n=3 B-Asns mice). **D.** Quantification of mitochondrial count as in Fig. 7G. Each data point represents a cell pooled from n=3 B-WT or B-Asns mice. Representative of two independent experiments. Statistical significance was determined by two-way ANOVA with Šidák’s (A-B,D) or with Tukey’s (C) multiple testing correction. Data are presented as the mean +/− SEM.

**Supplementary Figure 10.**
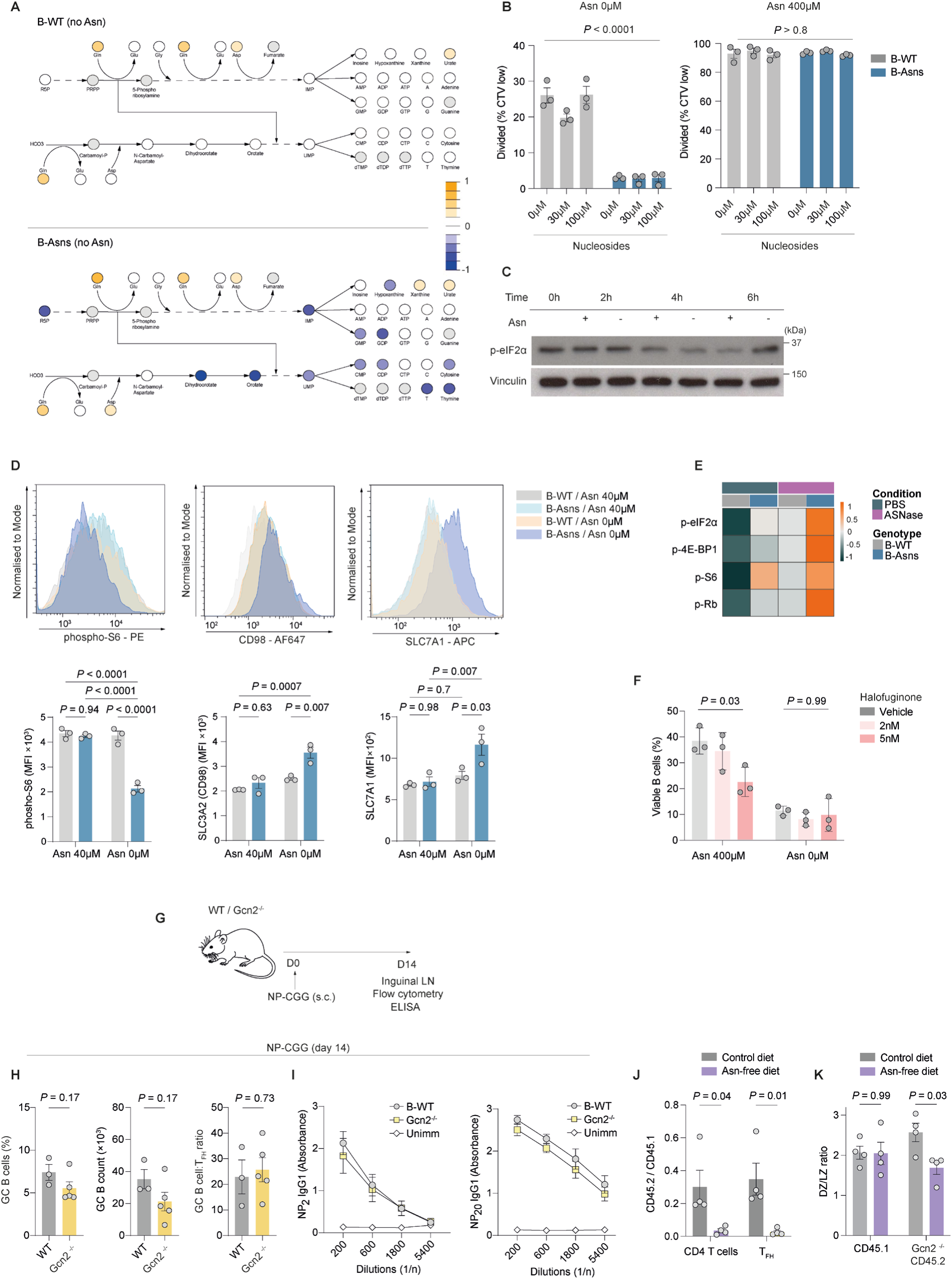
**A.** Relative abundance of metabolites in nucleotide synthesis pathway in B-WT and B-Asns B cells deprived of Asn. Grey-filled circles indicate that the metabolite was not detected. Scale represents abundance ratio. **B.** Quantification of the fraction of B cells from B-Asns and B-WT mice (n=3 mice) having diluting Celltrace Violet after stimulation for 72h with IL-4 and anti-CD40 in the presence or absence of Asn supplemented with adenosine, thymidine, uridine, cytidine, and guanosine at 0μM, 30μM or 100μM. Each data point represents a mouse. Data pooled from two independent experiments. **C.** Representative immunoblot time course of phospho-eIF2ɑ in wild type B cells stimulated with IL-4 and agonistic anti-CD40 in the absence or presence of Asn (400μM). Representative of two independent experiments. **D.** Representative flow plots and MFI quantifications of phospho-S6, SLC3A2 and SLC7A1 in B-WT or B-Asns B cells stimulated with IL-4 and anti-CD40 with 40μM Asn for 72h, then restimulated for an additional 24h in the presence (40μM) or absence of Asn (n=3 mice). Data pooled from two independent experiments. **E.** Heatmap of row z-scores for the gMFI values of phospho-EIF2α, phospho-4EBP1, and phosphoS6 and positive frequency of phospho-Rb in CD38^−^ GL-7^+^ GC B cells, measured by flow cytometry at day 9 SRBC immunisation following post-GC ASNase regime (mean of n = 4-5 mice each condition). Data pooled from two independent experiments **F.** Viability of wild type B cells stimulated with IL-4 and agonistic anti-CD40 for 72h, in the presence or absence of Asn (400μM), and with the indicated concentration of halofuginone or vehicle (DMSO). Representative of two independent experiments. **G.** Schematic of NP-CGG immunisation in Gcn2^−/−^ mice. WT and Gcn2^−/−^ mice were subcutaneously immunised with NP-CGG (50µg, on each flank) precipitated in Alum (1:1). On day 14, draining lymph node and serum were collected for analysis. **H.** Quantification of GC B cell proportions, absolute counts and relative counts to T_FH_ cells at day 14 post-immunisation. Data pooled from two independent experiments. **I.** Dilution curves of IgG1 and IgM anti-NP antibodies (NP_>2_-BSA and NP_20_-BSA respectively) at day 14, n=3-5 mice each condition. Data pooled from two independent experiments. **J.** CD45.2 Gcn2^−/−^ CD3^+^ CD4^+^ T cell and CD4^+^ BCL-6^+^ T_FH_ cell abundances relative to congenic CD45.1 WT counterparts under control or Asn-free diet setting. Data pooled from two independent experiments. **K.** DZ/LZ ratio quantification of CD45.2 Gcn2^−/−^ and congenic CD45.1 WT GC B cells (CD38^−^ BCL-6^+^) from bone marrow chimera experiment with two diet setting. Data pooled from two independent experiments. Statistical significance was determined by two-way ANOVA with Šidák’s (B,J,K) or Tukey’s (D,F) multiple testing correction, or unpaired two-tailed t test (H). Data are presented as the mean +/− SEM.

## Material and methods

### Study design

Statistical methods were not used to pre-determine sample sizes, but sample sizes were chosen to be comparable to those reported in previous studies. Effect sizes for certain experiments were estimated through pilot studies. Data distribution was assessed using normality tests to guide the selection of appropriate statistical methods, or it was assumed to be normally distributed. Mice that lacked germinal centres in the absence of Alum spots post-immunisation were considered to have failed intraperitoneal immunisation and were excluded from analysis. Biological and technical replicates were included in all experiments, with each experiment reflecting at least two independent replicates. All experiments included in the manuscript were found to be reproducible. For in vivo experiments, sex and age were matched in experimental batches. Efforts were made to minimise potential cage and litter effects by co-housing control and experimental mice, and using littermate controls where possible, although potential cage effects could not be avoided in diet experiments. Randomisation measures applied in in vivo experiments were also carried over into in vitro and ex vivo experiments, which were conducted with cells isolated from wild-type or knockout mice. During sample acquisition, experimental and control samples were run consecutively in an alternating fashion. Other randomisation methods were not applied, as they were not relevant to the study. For some experiments, researchers were blinded, such as when mice were genotyped after the experiment. However, data collection and analysis were not blinded in most experiments because the same researchers conducted the experiments and needed genotype information to ensure the inclusion of both wild-type and knockout mice in the study.

### Mice

C57BL/6J mice were obtained from Envigo. C57BL/6N-*Asns^tm1c(EUCOMM)Wtsi^*/H (EM:05307) mice were obtained from the Mary Lyon Centre, MRC Harwell, UK. B6.129S6-*Eif2ak4^tm1.2Dron^*/J (*Gcn2*^−/−^)(JAX: 008240), B6.C(Cg)-*Cd79a^tm1(cre)Reth^*/EhobJ (JAX: 020505), B6.129P2-*Aicda^tm1(Cre)Mnz^*/J (JAX: 007770), and B6;129S6-*Gt(ROSA)26Sor^tm9(CAG-tdTomato)Hze^*^/J^ (JAX: 007905) mice were obtained from Jackson Laboratories. B6.SJL.CD45.1 mice were provided by the central breeding facility of the University of Oxford. Male and female mice between the ages of 6-15 weeks were used. *Asns* ^LoxP/+^ × *Cd79a*-Cre^+/−^, *Asns^+/+^* × *Cd79a*-Cre^+/−^, *Asns* ^LoxP/+^ × *Cd79a*-Cre^−/−^, and *Asns* ^LoxP/ LoxP^ × *Cd79a*-Cre^−/−^ mice were used as controls (B-WT). All phenotypes were reproduced with all control genotypes.

Mice were bred and maintained under specific pathogen-free conditions at the Kennedy Institute of Rheumatology, University of Oxford. They were housed in cages that had individual ventilation and were provided with environmental enrichment. The temperature was kept between 20-24°C, with a humidity level of 45-65%. They were exposed to a 12-hour cycle of light and darkness (7 am to 7 pm), with a thirty-minute period of dawn and dusk. All procedures and experiments were performed in accordance with the UK Scientific Procedures Act (1986) under a project license authorized by the UK Home Office (PPL number: PP1971784).

### *In vivo* modification of asparagine levels

Experimental and control mice were intraperitoneally injected with ≈10U (500U/kg) of asparaginase from *E. coli* (Abcam, ab277068) diluted in PBS, according to experimental regime. Two main regimes were used for ASNase experiments: ‘standard’ and ‘post-GC’. In the standard schedule, ASNase was administered from one day prior to immunisation, and then every two days over a nine day period before analysis. In the post-GC schedule, only two doses of ASNase were given, at days six and eight after SRBC-immunisation or at days 6, 9 and 12 after NP-CGG immunisation, thereby targeting established GCs.

Asn-free diet or control chow (Research Diets, A05080216i and A10021Bi) was administered for 12 days before or 7 days after immunogenic challenges and maintained for the duration of the experiments. The detailed formula of diet amino acid constituents is indicated below.

**Table:**
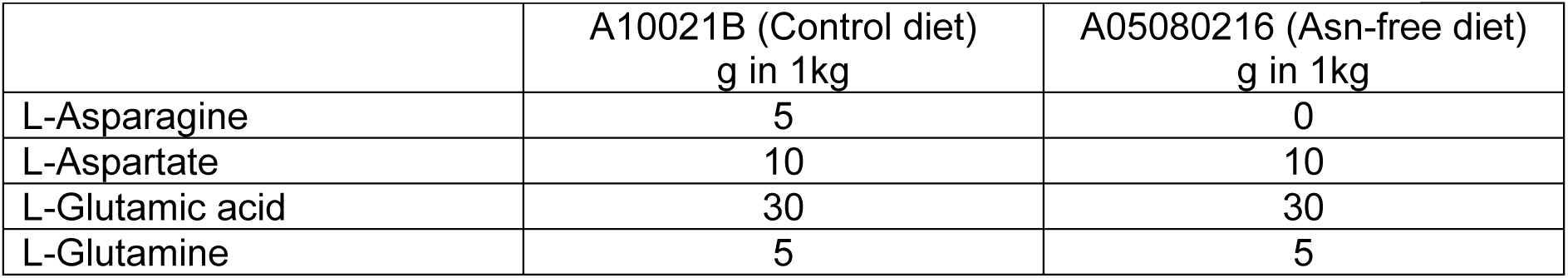
Diet amino acid constituents provided by the vendor (Research Diets)

|  | A10021B (Control diet)<br>g in 1kg | A05080216 (Asn-free diet)<br>g in 1kg |
| --- | --- | --- |
| L-Asparagine | 5 | 0 |
| L-Aspartate | 10 | 10 |
| L-Glutamic acid | 30 | 30 |
| L-Glutamine | 5 | 5 |

Drinking water was supplemented with L-asparagine monohydrate (Merck, cat: A7094-25G) at 1.5 g/L and administered to mice two weeks prior to immunisation and maintained for the whole duration of the experiment. Drinking water was replaced with freshly prepared Asn every 2-3 days.

### Immunisation

One ml of sterile SRBCs in Alsever’s solution (ThermoFisher, EO Labs, or TCS bioscience) were washed twice with 10ml of ice-cold PBS and reconstituted in 2-3ml of PBS, and 200μl injected intraperitoneally or intravenously. In some experiments, an enhanced SRBC immunisation method was used to maximise GC B cell yield, by immunising mice with 0.1ml SRBC on day 0 followed by a second injection of 0.2ml on day 4 or 5^53^. For protein antigen immunisations, 50μg NP_(30-39)_-CGG (Biosearch Tech, cat: N-5055D-5) in PBS was mixed with Imject Alum (ThermoFisher) or Alum Hydrogel (InvivoGen, cat: vac-alu-50) at a 1:1 ratio (vol:vol) and rotated at room temperature for 30 mins (used for Imject Alum) before intraperitoneal injection in 100μl volume, or mixed vigorously with a pipette for 5 mins (used for Alum Hydrogel) before subcutaneous injection in the flanks, hock, or intraperitoneal space. 50μg NP-AECM-FICOLL (Biosearch) in PBS was mixed with PBS at a 1:1 ratio and intraperitoneally injected.

### Influenza infection

The A/HK-X31 (X31, H3N2) strain was used. Mice were anaesthetised using isoflurane and intranasally inoculated with 5×10^3^ PFU of X31 influenza A virus in PBS. Mice were closely monitored and weighed for 14 days following infection, and characteristic weight loss was confirmed in all infected mice.

### Bone marrow chimera generation

B6.SJL.CD45.1 recipient mice were administered two doses of 5.5Gy irradiation four hours apart. Mice were then intravenously injected with 4×10^6^ mixed bone marrow (BM) cells at a 1:1 ratio, isolated from age- and sex-matched CD45.2^+^ B-WT or B-Asns or *Gcn2*^−/−^ mice, and CD45.1^+^ WT donor mice. Recipient mice were maintained on antibiotics (Baytril, Bayer corporation) administered in their drinking water for two weeks. Bone marrow reconstitution was confirmed by flow cytometry of peripheral blood at 8 weeks. Asn-free diet or control chow (Research Diets, A05080216i and A10021Bi) was administered for 12 days starting from week 8, maintained for additional 9 days (total of 21 days) during immunisation with SRBC until mice were terminated at 11 weeks.

### Cell isolation

Spleens were dissociated by passing through 70μm cell strainers. For *ex vivo* GC B cell mass spectrometry, GC B cells were first enriched using the mouse Germinal Center B Cell (PNA) MicroBead Kit (Miltenyi), and then further purified by flow sorting (Live/Dead^−^ CD19^+^IgD^−^GL-7^+^CD95^+^). Total B cells from the same mouse pool were pre-enriched using CD19^+^ Microbeads (Miltenyi), then naïve B cells purified by flow sorting (Live/Dead^−^ B220^+^IgD^+^).

To isolate cells for RNA extraction, GC B cells (Live/Dead^−^CD19^+^IgD^−^CD95^+^GL-7^+^) and naïve B cells (Live/Dead^−^CD19^+^IgD^+^) were flow sorted into RLT Plus buffer (Qiagen) following pre-enrichment with the Pan B Cell Isolation Kit II (Miltenyi). B cells for culture were isolated using the Pan B Cell Isolation Kit II (Miltenyi). Purity was routinely >90% by flow cytometry.

For some experiments, untouched GC B cells were isolated using a magnetic bead-based protocol as described^54^. Briefly, single cell suspensions were prepared from spleens of SRBC-immunised mice (enhanced protocol) or NP-CGG-immunised mice (scRNAseq experiment) in ice cold MACS isolation buffer (PBS with 0.5% BSA and 2mM EDTA) followed by ACK (Gibco) RBC lysis for 4 mins at 20°C. Following washing, cells were labelled with anti-CD43 microbeads (Miltenyi) and biotinylated antibodies against CD38 and CD11c (both eBioscience, clones 90 and N418 respectively) and IgD (Thermofisher, clone 11-26c.2a). Then, cells were incubated with anti-biotin Microbeads (Miltenyi), and subsequently run through an LS column (Miltenyi). GC B cell purity was typically around 90%. For scRNAseq, GC B cells were further purified using fluorescence-activated cell sorting (FACS).

### Cell culture

Total B cells were isolated as described above. B cells were cultured at 1-3×10^6^ cells/ml in RPMI1640 (custom product from Cell Culture Technologies, Gravesano, Switzerland, lacking glutamate, aspartate, glutamine, and asparagine), supplemented with 1mM pyruvate, 10mM HEPES, 100 IU/ml penicillin/streptomycin and 50μM 2-mercaptoethanol, and 10% dialysed FBS (Gibco), Aspartate (Asp) (150μM), Glutamate (Glu) (140μM), GlutaMAX (ThermoFisher or Sigma) (2mM), indicated concentration of Asn, except where mentioned otherwise. B cells were stimulated with agonistic anti-CD40 (5μg/ml, FGK45.4, functional grade, Miltenyi), recombinant IL-4 (1,10 or 50ng/ml, Peprotech), LPS (10μg/ml, Merck) or CpG (100nM, ODN1826, Miltenyi).

For proliferation experiments, cells were labelled with CellTrace Violet (ThermoFisher) according to manufacturer’s instructions before stimulation. For stable isotope labelling, ^15^N(amide)-asparagine (485896, Merck) (400μM), ^15^N(amide)-glutamine (NLM-557, Cambridge Isotope Laboratories) (2mM), ^15^N(amino)-glutamine (NLM-1016, Cambridge Isotope Laboratories) (2mM), or ^13^C_5_-glutamine (CLM-1822, Cambridge Isotope Laboratories) (2mM) were used, replacing their unlabelled molecule in RPMI1640.

For nucleoside rescue experiment, adenosine (Fluorochem cat: F093333), thymidine (Fluorochem cat: F078885), uridine (Fluorochem cat: F078878), cytidine (Fluorochem cat: F226675) and guanosine (Merck, cat: G6264-5G) were prepared freshly in water and added to culture media at a final concentration of 30μM or 100μM.

Halofuginone hydrobromide (Bio-techne, cat no: 1993) was reconstituted in DMSO and used at 2nM and 5nM in the culture. Dimethyl 2-oxoglutarate (DM-OG) was purchased from Merck (cat no: 349631-5G) and added to the culture medium at 2mM and 4mM final concentrations.

### *Ex vivo* lymph node slice culture

Inguinal lymph nodes (LNs) were harvested from B-WT or B-Asns mice immunised with NP-CGG (precipitated in Alum hydrogel) subcutaneously on both flanks. They were then removed from surrounding fat by dissection and mild detergent wash (<1 sec) in 0.01% digitonin solution (Thermo Scientific) and embedded in 6% w/v low melting point agarose (Lonza) in 1x PBS. 300μm thick slices were cut using the Precisionary Compresstome® VF-210-0Z following manufacturer’s instructions with a blade speed of 4 and oscillation of 6. The buffer tank was filled with ice cold PBS containing 1% Pen/Strep. Slices were removed from surrounding agarose and placed in a 24 well plate in 500μl of Asn-replete or Asn-free complete RPMI media in duplicate. LN slices were allowed to rest and equilibrate in the media for 1 hour at 37°C and 5% CO2 before transferral into fresh media and incubation for 20hrs. Following culture, slices were collected, duplicate slices from same lymph nodes pooled, and slices disrupted into PBS containing 2% FBS and 2mM EDTA using the tip of an insulin syringe plunger to generate a cell suspension in a 96-well U bottom plate. Cells were then stained with viability dye and surface antigens. Following washes, cells were fixed in fresh 4% PFA (Cell Signalling) for 15 mins at 20°C and permeabilised in methanol for intracellular staining and subsequently run on an Aurora flow cytometer (Cytek).

### Flow cytometry

Spleens were dissociated by passing through 70μm cell strainers, and red cells lysed with ACK Lysis Buffer (Gibco). Single cell suspensions were incubated with Fixable Viability Dye eFluor™ 780 (eBioscience) or Zombie NIR (BioLegend) in PBS, supplemented with Fc Block and then surface antibodies (30 mins on ice) in FACS buffer (PBS supplemented with 0.5% BSA and 2mM EDTA). For intracellular staining, cells were fixed in 4% paraformaldehyde at 20°C, then permeabilised with ice cold 90% methanol for 10 mins. For FoxP3 staining, eBioscience™ Foxp3 / Transcription Factor Staining Buffer Set (ThermoFisher) was used. To measure OPP ex vivo incorporation, cells were incubated in RPMI1640 with the indicated concentration of Asn, in the presence of 20μM OPP for 25 mins at 37°C. Click labelling was then performed with the Click-iT™ Plus OPP Alexa Fluor 647 Protein Synthesis Assay Kit (ThermoFisher, cat: C10458). Apoptosis was quantified by annexin V-FITC binding (Biolegend). The proliferation index was determined using the Proliferation module of FlowJo (BD). SRBC binding assay was performed as previously reported^55^. Briefly, splenocytes were harvested from mice immunised with SRBC. Following surface staining, cells were mixed with SRBCs labelled with 10µM Cell Trace Violet (ThermoFisher) on ice for 20 mins. After washing, cells were resuspended in FACS buffer and acquired on BD Fortessa X20.

| **Antibody** (Catalogue number) | **Clone** | **Manufacturer** |
| --- | --- | --- |
| Anti-GL-7 (144612) | GL-7 | Biolegend |
| Anti-CD138(142504) | 281-2 | Biolegend |
| Anti-CD19 (115538) | 6D5 | Biolegend |
| Anti-IgD (405721) | 11-26c-2a | Biolegend |
| Anti-CD38 (102718) | 90 | Biolegend |
| Anti-CXCR4 (146513) | L276F12 | Biolegend |
| Anti-CD98 (128211) | RL388 | Biolegend |
| Anti-IgG1 (406617) | RMG1-1 | Biolegend |
| Anti-MHCII (107605) | M5/114.15.2 | Biolegend |
| Anti-CD23 (101608) | B3B4 | Biolegend |
| Anti-CD43 (12-0431-82) | eBioR2/60 | ThermoFisher |
| Anti-CD86 (105031) | GL-1 | Biolegend |
| Anti-B220 (103247) | RA3-6B2 | Biolegend |
| Anti-IgM (406513) | RMM-1 | Biolegend |
| Anti-CD21/35 (123409) | 7E9 | Biolegend |
| Anti-TACI (133403) | 8F10 | Biolegend |
| Anti-CD16/32 (Fc Block) | 2.4G2 | ThermoFisher |
| Anti-CXCR4 (741783) | 2B11 | BD Bioscience |
| Anti-CD38 (741955) | 90 | BD Bioscience |
| Anti-CD4 (100547) | RM4-5 | Biolegend |
| Anti-PD-1 (135213) | 29F.1A12 | Biolegend |
| Anti-CXCR5 (145511) | L138D7 | Biolegend |
| Anti-CD45.1 (110717) | A20 | Biolegend |
| Anti-CD45.2 (109817) | 104 | Biolegend |
| Anti-CD3 (100215) | 17A2 | Biolegend |
| Anti-IRF4 (12-9858-82) | 3E4 | ThermoFisher |
| Anti-BCL6 (561525) | K112-91 | BD Bioscience |
| Anti-CHOP (NB600-1335AF700) | 9C8 | Novus Bio |
| Anti-Foxp3 (25-5773-82) | FJK-16 | ThermoFisher |
| Anti-phospho-Rb (Ser807/811) (8516T) | D20B12 | Cell Signalling Technologies |
| Anti-phospho-S6 (Cat #17-9007-42) | cupk43k | ThermoFisher |
| Anti-phospho-EIF2α (EIF2S1) (CL488-68023) | 1A4A11 | Proteintech |
| Anti-phospho-4EBP1 (12-9107-42) | V3NTY24 | ThermoFisher |
| Anti-COXIV (ab197491) | mAbcam33985 | Abcam |
| Anti-COXIV (CL488-60251) | 3C7D2 | Proteintech |
| Anti-COX I (MTCO1) (ab198600 & ab154477) | 1D6E1A8 | Abcam |
| Anti-Cytochrome C (NB100-56503AF700) | 7H8.2C12 | Novus |
| Anti-SDHA (ab170172) | 2E3GC12FB2AE2 | Abcam |
| Anti-Cyclin B1 (647903) | V152 | Biolegend |
| Anti-phospho-H3 (3465S) | D2C8 | Cell Signalling Technologies |

### *Ex vivo* single cell competitive Asn uptake assay

We followed the recently published QUAS-R protocol with certain modifications^27^. Briefly, spleens were harvested 8 days post SRBC immunisation and dissociated through 70μm cell strainers. Red blood cells were lysed with ACK lysis buffer and single cell suspensions were incubated with Zombie NIR (BioLegend) and Fc block in PBS on ice for 30 minutes followed by surface staining in FACS buffer for 30 minutes on ice. Hanks’ Balanced Salt Solution HBSS (ThermoFisher) supplemented with Click-iT homopropargylglycine HPG (ThermoFisher) at 400μM and L-asparagine monohydrate (Merck) at increasing concentrations (0, 10, 40, 100, 400, 4000μM) were warmed at 37°C. Cells were resuspended in HBSS and 50μl of suspension was plated and incubated at 37°C for 10 minutes. Control cells incubated at 4°C were kept on ice for background detection. HPG-containing media supplemented with each Asn dilution was added and cells were further incubated for 3 minutes, before immediate fixation with PFA at 4% for 25 minutes at room temperature. Cells were washed and permeabilised with 1× saponin-based permeabilization buffer (ThermoFisher) for 20 minutes at room temperature before adding the click chemistry reaction mix, using the Click-iT™ Plus OPP Alexa Fluor™ 647 Protein Synthesis kit due to the shared Click Chemistry of HPG with OPP and incubating for 30 minutes at room temperature. Cells were washed and resuspended in FACS buffer for flow cytometry acquisition.

### Immunofluorescence microscopy and image analysis

Spleens were fixed overnight in Antigenfix (DiaPath) solution at 4°C. The next day, spleens were washed in PBS, followed by overnight incubation in 30% sucrose (in PBS) at 4°C for cryoprotection. On the following day, spleens were snap frozen in 100% methanol on dry ice and stored at −80°C until cryosectioning at 8μm thickness. Slides were then rehydrated in PBS at 20°C, then blocked in PBS containing 0.05% Tween-20, 10% goat serum, and 10% rat serum at RT for one hour. All primary antibody staining was performed overnight at 4°C or 1 hour at RT in PBS supplemented with 2% rat serum and 0.05% Tween-20. For visualising GCs and B cell follicles, anti-GL-7 AF488 (Biolegend, clone: GL7) and anti-CD21/35 (Biolegend clone: 7E9) were used at 1:50 and 1:100 dilutions, respectively.

To visualise protein synthesis *in vivo*, mice were immunised with SRBC using the enhanced protocol. At day 14, they were injected intraperitoneally with 1.5mg L-AHA (Thermo) and then 4 hours later 1mg OPP (Jena Bioscience) dissolved in 100μl PBS, or PBS alone as a negative control. Mice were sacrificed 1h later, and spleens were harvested for immunofluorescence analysis. L-AHA and OPP were detected by Click Chemistry using the Click-iT™ Plus L-AHA and then Click-iT™ Plus OPP Alexa Fluor™ 488 or 647 Protein Synthesis kits (Thermo) following the manufacturer’s protocols.

Slides were mounted with Fluoromount G (Southern Biotech, cat: 0100-01) and imaged using a Zeiss LSM 980 equipped with an Airyscan 2 module or Zeiss Axioscan 7.

Image analysis was performed in ImageJ. After adjusting brightness and contrast automatically, GCs were identified as GL-7^+^ clusters and manually counted on each splenic section. For area quantification, GCs were selected either manually or using autothresholding (Yen) and registered as Region of interests (ROIs). Then, area was quantified using Measure or Analyze particles function with 750×m^2^ threshold.

### Multiplexed Tissue Imaging using Cell DIVE

Formalin-fixed paraffin embedded (FFPE) human tonsil sections (5μm) were deparaffinised and rehydrated, and then permeabilised in 0.3% Triton X100 for 10 mins followed by washing in PBS for 5 mins. Antigen retrieval was performed in a Biocare NxGen decloaker by boiling the slides in a citrate-based antigen unmasking solution (Vector, #H3300) at pH 6, and then in a Tris-based antigen retrieval solution, containing 1×Tris, EDTA, 0.05% Tween20 and ddH2O (pH 9), for 20 mins at each step. The slides were blocked overnight in PBS solution containing 3% BSA (Merck, #A7906), 10% donkey serum (Bio-Rad, #C06SB), then incubated with human FcR Blocking Reagent (Miltenyi) at 1:100 at room temperature for 1 hour. After washing in PBS, the slides were stained with DAPI (ThermoFisher), washed again, and cover slips were placed with mounting media (50% glycerol, [Sigma] and 4% propyl gallate [Sigma, #2370]).

All FFPE human tonsil slides were imaged using a GE Cell DIVE system. Firstly, a ScanPlan was performed by acquiring at 10× magnification so that regions of interest could be selected. This was then followed by imaging at 20× to acquire background autofluorescence and generate virtual H&E images. This background was subtracted from all subsequent rounds of staining.

All slides were decoverslipped in PBS before each staining round, which consisted of three antibodies applied in staining buffer (3% BSA in PBS). Unconjugated primary antibodies were used in the first round, incubated at 4°C overnight. The next day, slides were washed in 0.05% PBS-Tween 20, (Sigma, P9416) three times. AlexaFluor 488, 555 or 647 conjugated secondary antibodies were then added to the slides for an hour at room temperature. Directly conjugated antibodies were used in subsequent staining rounds, incubated at 4°C overnight. Antibody conjugation was performed in house for some of the antibodies using the antibody labelling kit (Invitrogen).

Fluorescent dye inactivation was performed by bleaching the slides with 0.1 M NaHCO3 (pH 10.9-11.3) (Sigma, S6297) and 3% H_2_O_2_ (Merck, #216763,) for 3 rounds (15 mins each) with a 1 min wash in PBS in between. The slides were then re-stained with DAPI for 2 mins before being washed in PBS. The bleached slides were imaged as the new background for subsequent subtraction.

Image analysis was performed in ImageJ. 225μm^2^ region of interests (ROIs) were selected randomly in CD21^−^ IgD^−^ DZ GC, CD21^+^ IgD^−^ LZ GC and surrounding IgD^+^ B cell follicles and CD3^+^ T cell areas of individual GCs with representative DZ-LZ distribution. These ROIs were then inspected in the ASNS channel and mean fluorescence was calculated using the Measure function. Seven GCs from each tonsil were pooled for analysis, and a total of 21 GCs from 3 tonsils were quantified.

#### Antibody list

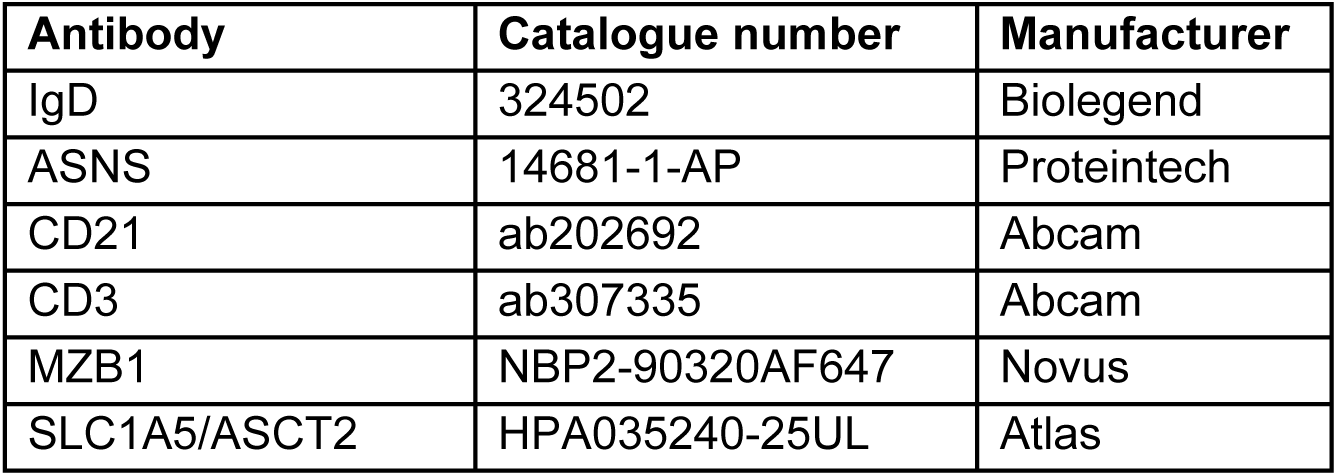

### 3D Lattice SIM

Cultured B cells were transferred onto poly-L-lysine coated coverslips and incubated at 37°C for 10 minutes. Cells were subsequently fixed in 4% paraformaldehyde for 10 minutes at 37°C, before permeabilisation and blocking at 20°C for 30 minutes in PBS containing 0.2% Triton X-100, 10% goat serum and 10% rat serum. Staining with conjugated antibodies was then performed overnight at 4°C in PBS supplemented with 2% goat serum. Nuclear staining was carried out with DAPI (Sigma) diluted in PBS for 15 minutes at 20°C, prior to coverslip mounting with Fluoromount G (Southern Biotech, catalogue no. 0100-01). Imaging was performed with the Zeiss Elyra 7 lattice SIM using the 63× oil immersion objective and 15 phase SIM. SIM processing was then carried out on ZEN Black.

Image quantification of the mitochondrial network was performed using ImageJ software. Single cells were defined in individual Region of Interest (ROI) based on COX I signal. 3D mitochondrial morphological and network analysis was subsequently performed using an ImageJ plugin^56^ with modified parameters: rolling (microns) 1.28 and gamma (0.90).

### Extracellular flux analysis (SeaHorse)

Real-time oxygen consumption rates (OCR) and extracellular acidification rates (ECAR) were measured using a Seahorse XFe96 Extracellular Flux Analyzer (Agilent). Ex vivo isolated B cells from B-WT and B-Asns mice were stimulated (anti-CD40 at 5μg/ml and IL-4 at 10ng/ml) overnight at 10^6^ cells per ml in a 24 well plate in the presence of varying concentrations of Asn. The next day, 2×10^5^ B cells were plated on a poly-D lysine (Sigma)-coated XF96 cell culture microplate with 5-6 technical replicate wells and incubated at 37 °C for a minimum of 30 min in a CO_2_-free incubator in assay medium (XF DMEM medium pH 7.40 supplemented with 2 mM L-glutamine, 1 mM pyruvate and 10 mM Glucose) before the initiation of the measurements. Basal OCR and ECAR were measured, then followed by the MitoStress test, with oligomycin (1μM), fluorocarbonyl cyanide phenylhydrazone (FCCP, 2μM) and rotenone + antimycin A (0.5μM) (Agilent). Analyses were performed on Wave (Agilent).

### ELISA

96 well EIA/RIA plates (Corning, cat: 3590) were coated with NP conjugated with BSA (NP_2_ or NP_20>_, cat: N5050XL-10 and cat: N-5050H-10, both from BioSearch) at 5μg/ml in bicarbonate/carbonate coating buffer overnight at 4°C. The next day, plates were washed with PBS and blocked with 5% skimmed milk in PBS for 2-2.5 hours at 37°C. Sera obtained from mice (NP-CGG-immunisation day 14 subcutaneously or unimmunized control) were serially diluted in 1% skimmed milk and added into blocked plates for exactly 1hr at 37°C. After several washes with PBS-Tween20 (0.05%), alkaline phosphatase-conjugated goat anti-mouse IgG1 (Southern Biotech, cat: 1071-04) or IgM (Southern Biotech, cat: 1021-04) detection antibodies (at 1:2000 dilution) were incubated for exactly 1hr at 37°C. After the final washing step, plates were developed with alkaline phosphatase substrate (Sigma, cat: P7998-100ML) with simultaneous reading on FLUOstar Omega plate reader every 5 mins.

96 well EIA/RIA plates were coated overnight at 4°C with PFA-inactivated X31 virus in bicarbonate/carbonate coating buffer. After blocking with 1% BSA in PBS-Tween (0.05%) at 20°C for 2h, serially-diluted sera added and incubated further 2h at 20°C. After multiple washes, biotinylated goat anti-mouse IgG or IgM detection antibodies (Southern Biotech, 1:1000) were added and incubated at 20°C for 1h. Following washes, Horseradish Peroxidase (HRP) conjugated with Streptavidin (Jackson ImmunoResearch, 1:1000) was added and incubated for additional 30 mins. The 3,3′,5,5′-Tetramethylbenzidine (TMB substrate) (ThermoFisher) added following multiple washes. Reading was performed at two wavelengths (450 and 570) with precise scanning mode after stopping the reaction with 2N H_2_SO_4_.

For avidity assessment, following serum incubation, there was an additional 15-minute washing step at 20°C with freshly prepared urea (vWr) in PBS at concentrations of 1, 2.5, or 5M. The detection and development procedures remained the same as the aforementioned protocol.

### Serum and lymph node interstitial fluid (LNIF) collection for LC-MS

For serum collection, mice were either bled from the tail vein or subjected to terminal bleeding through intracardiac procedures performed after CO₂ asphyxiation. The collected blood was then centrifuged at 1000 × g for 10 minutes at 4°C, and the serum was stored at −20°C until analysis.

For LNIF collection, mice were humanely euthanized using CO₂ asphyxiation, and the total inguinal, brachial, axial, cervical, mediastinal, and mesenteric lymph nodes were collected and placed in 1.5ml tubes filled with PBS, and placed on ice. After harvesting lymph nodes from 9-14 mice into two to three 1.5ml tubes, the elution phase was commenced. Lymph nodes from four mice were pooled in individual tubes, treated as a batch, and processed together for lymph node interstitial fluid (LNIF) elution. Lymph nodes pooled from four mice in a single Eppendorf tube were quickly dabbed on Kimtech paper (Kimberly-Clark), separated into two groups, and placed on two 10µm strainers (pluriSelect Cat: 43-10010-40) inserted into 1.5ml tubes. These were then centrifuged at 400 × g for 10 minutes at 4°C. Following the first spin, the strainers were reversed and centrifuged again at 400 × g for another 10 minutes at 4°C to maximize fluid output.

The LNIF collected at the bottom of the tube was gently transferred to another collection tube using a pipette, carefully avoiding any contaminating cell pellets if present. The LNIF was pooled with that from the subsequent tubes/batches. The final pooled volume from 9-14 mice was treated as one biological replicate, and the procedure was repeated two more times to obtain three biological replicates (a total of 35 mice were used, excluding those used for serum isolation). Using this method, it was possible to collect approximately 1-1.5µl of LNIF per mouse.

### Mass spectrometry

To extract molecules, cell were pelleted and washed twice with cold PBS and once with ice cold ammonium acetate (150mM, pH 7.3). The pellet was then resuspended in 1ml 80% LC-MS grade methanol/ultrapure water chilled on dry ice. The samples were vortexed three times on dry ice, and then centrifuged at 15,000×g for 5 mins. The initial supernatant was then transferred into a 1.8ml glass vial, and the pellet resuspended in 200μl 80% methanol as above, then centrifuged again at 15,000×g for 5 mins. The second supernatant was then combined with the first, dried at 4°C using a Centrivap (Conco), then stored at −80°C until analysis.

Dried metabolites were resuspended in 50% ACN:water and 1/10^th^ was loaded onto a Luna 3μm NH_2_ 100A (150 × 2.0 mm) column (Phenomenex). The chromatographic separation was performed on a Vanquish Flex (Thermo Scientific) with mobile phases A (5mM NH_4_AcO pH 9.9) and B (ACN) and a flow rate of 200μl/min. A linear gradient from 15% A to 95% A over 18 min was followed by 7 min isocratic flow at 95% A and reequilibration to 15% A. Metabolites were detected with a Thermo Scientific Q Exactive mass spectrometer run with polarity switching (+3.5 kV/− 3.5 kV) in full scan mode with an m/z range of 70-975 and 140.000 resolution. Maven (v8.1.27.11) was used to quantify the targeted metabolites by area under the curve using expected retention time (as determined with pure standards) and accurate mass measurements (< 5 ppm). Values were normalized to cell number.

Relative amounts of metabolites were calculated by summing up the values for all measured isotopologues of the targeted metabolites. Metabolite Isotopologue Distributions were corrected for natural ^15^N abundance. Integrated pathway analysis was performed using MetaboAnalyst^57^ (https://www.metaboanalyst.ca/) with default parameters. This function uses over-representation analysis (ORA) based on the hypergeometric distribution, to generate *P* values for the over representation of genes or metabolites within KEGG pathways. The *P* values are then merged, and network topology analysis used to determine an impact score for the pathway. The input metabolite and gene sets were those differentially expressed/abundant in GC B cells compared with follicular B cells (using multiple testing correction).

For amino acid detection in serum and lymph node intersistial fluid following LC-MS protocol was used. Data acquisition was performed using an adaptation of a method previously described^58^. Samples were injected onto a Dionex UltiMate 3000 LC system (Thermo Scientific) with a Fortis C18 column (100 x 2.1 mm, 3 μm; Fortis Technology Ltd). Mobile phases: Solvent A was 0.1 % acetic acid in water (both Optima HPLC grade, Sigma Aldrich); solvent B was Acetonitrile (Optima HPLC grade, Sigma Aldrich). The column temperature, flow rate, and injection volume were 50 °C, 0.2 mL min-1, and 5 μL, respectively. Gradient elution was as follows: 0% B, 0-3 min; 0 – 60% B, 3-5 min; hold for 2 min; 0% B, 7.01, hold for 4 min (re-equilibration). MS was performed in positive and negative polarities using a TQS Quantiva Triple Quadrupole Mass Spectrometer (Thermo Scientific) with an electrospray ionization (ESI) source. Qualitative and quantitative analyses were performed using Xcalibur Qual Browser and Tracefinder 5.1 software (Thermo Scientific) according to the manufacturer’s workflows. Analyses were performed using selected reaction monitoring (SRM). Precursor-to-product ion transitions and collision energies are listed in Table 1 below. Results were processed using GraphPad Prism 10 software.

**Table:** SRM parameters.

| Compound | RT (min) | Precursor (m/z) | Product (m/z) | Collision Energy (V) | Ionization |
| --- | --- | --- | --- | --- | --- |
| Asparagine | 1.11 | 133.12 | 74<br>87.058 | 16.67<br>10.25 | Positive |
| Glutamine | 1.15 | 147.138 | 84<br>130.058 | 17.69<br>10.25 | Positive |
| <sup>13</sup>C<sub>4</sub>-Asparagine | 1.11 | 137.132 | 90 | 10.25 | Positive |
| <sup>13</sup>C<sub>4</sub>-Asparagine | 1.11 | 135.22 | 117 | 10.25 | Negative |
| <sup>13</sup>C<sub>5</sub>-Glutamine | 1.15 | 152.152 | 135.04<br>88 | 10.25<br>17.84 | Positive |

Samples were prepared as follows: (a) 20 μL serum were mixed with 60 μL MeOH containing internal standard (IS, 100 μmol L-1 13C5-glutamine and 50 μmol L-1 of 13C4-asparagine), vortexed briefly and centrifugated (10 min, 4 °C, 12,700 rpm). 60 μL supernatant was collected and dried under a N2 stream. Samples were resuspended in 60 μL water, vortexed briefly, and analysed by LC-MS (see above). (b) 15 μL mouse lymph node interstitial fluid (LNIF) and mixed with 45 μL methanol containing IS (same concentration as above). Sample preparation and analysis continued as above, but with a final resuspension volume of 45 μL water. For both a and b relative abundances of metabolites were measured by comparison of their peak areas compared to those of their labelled equivalents (i.e. IS). For serum, calibration curves (300, 150, 75, 37.5, 18.8, 9.3, and 4.6 µmol L-1) for labelled IS were generated using a matrix of human serum, prepared as above.

Reagents were as follows: Optima water (Optima HPLC grade, Sigma Aldrich), Acetonitrile (Optima HPLC grade, Sigma Aldrich), Acetic acid (Fisher Chemical), Methanol (Optima HPLC grade, Sigma Aldrich), 13C4-asparagine (Cambridge Isotope Laboratories, Item number CLM-8699-H-PK), 13C5-glutamine (Cambridge Isotope Laboratories, C.N 184161-19-1).

### Western blotting

Cells were lysed on ice in RIPA buffer (Merck) supplemented with protease and phosphatase inhibitors (complete™ ULTRA Tablets and PhosSTOP™, Roche) for 30mins with frequent vortexing, and then centrifuged at 15,000×g for 15mins at 4°C and the supernatant recovered. Protein was quantified using the BCA method (Pierce, Thermo). Samples were denatured in Laemmli buffer (BioRad) containing 10% β-mercaptoethanol at 90°C for 5 mins then transferred to ice, before running on a 4-15% gel (Mini-PROTEAN TGX Precast Gels, BioRad). Protein was transferred onto PVDF membranes (BioRad), which were blocked in 2.5% skimmed milk for 1h, and then stained with the primary antibody diluted in 2.5% BSA/0.05% TBS-Tween 20 at 4°C overnight.

Following washing, membranes were then incubated with horseradish peroxidase (HRP)-conjugated goat anti-rabbit secondary antibody in 2.5% milk/0.05% TBS-Tween 20 for 1h at RT. Chemiluminescence was used to detect the secondary antibody (Anti-rabbit IgG, HRP-linked Antibody #7074 and SuperSignal™ West Pico PLUS Chemiluminescent Substrate, ThermoFisher).

| Antibody (Catalogue number) | Dilution | Clone | Manufacturer |
| --- | --- | --- | --- |
| DHODH (14877-1-AP) | 1:1000 | Polyclonal | Proteintech |
| ASNS (14681-1-AP) | 1:2000 | Polyclonal | Proteintech |
| CHOP (2895) | 1:1000 | L63F7 | Cell Signaling |
| Phospho-4E-BP1 (236B4) | 1:500 | Monoclonal | Cell Signaling |
| ATF4 (11815) | 1:500 | D4B8 | Cell Signaling |
| Vinculin (13901) | 1:1000 | E1E9V | Cell Signaling |
| β-actin (4970) | 1:2000 | 13E5 | Cell Signaling |
| ATIC (ab188321) | 1:500 | EPR13243-53 | Abcam |
| Phospho-eIF2α (9721) | 1:500 | Polyclonal | Cell Signaling |

### Quantitative PCR

Cells were washed in PBS and lysed in RLT Plus buffer (Qiagen) supplemented with 1% β-mercaptoethanol. RNA extraction was performed according to manufacturer’s instructions using an RNeasy Plus Mini Kit (Qiagen). Reverse transcription was performed using the High Capacity RNA-to-cDNA Kit (Thermo). Quantitative qPCR was performed using Taqman Probes (Thermo). Relative expression was calculated with reference to *Ubc or Hypoxanthine-guanine phosphoribosyltransferase (Hprt)*.

List of probes used:

| Gene | Taqman ProbeID |
| --- | --- |
| *Asns* | Mm00803785_m1 |
| *Ubc* | Mm02525934_g1 |
| *Atf4* | Mm00515325_g1 |
| *Got2* | Mm00494703_m1 |
| *Hprt* | Mm03024075_m1 |

### Single cell RNA sequencing

Following labelling with TotalSeq-C anti-mouse Hashtag oligonucleotide-conjugated antibodies (Biolegend) and flow sorting, GC B cells from individual mice were combined into three pools, each containing cells from one mouse of each experimental condition. The cell count and viability were determined with acridine orange/propidium iodide fluorescence using a LUNA-FX7 cell counter (Logos Biosystems). 10,000-15,000 cells per pool were loaded onto the 10X Genomics Chromium X instrument (Chip K). Gene expression and Hashtag oligonucleotide sequencing libraries were prepared using the 10x Genomics Single Cell 5’ Reagent Kits v2 (Dual Index) following manufacturer user guide (CG000330 Rev B). The final libraries were loaded on the NovaSeq X sequencing platform (Illumina, 150bp paired end).

Filtered matrices were generated using Cellranger 7.1 and pre-processed with *Scanpy*. Individual samples were demultiplexed and doublets removed using *Solo*. Cells with fewer than 300 genes, or genes found in fewer than 10 cells were removed, as were those with more than 7% mitochondrial genes. Immunoglobulin genes were removed. Sequencing batches were integrated using multiresolution variational inference, implemented using the *MrVI* model in *scVI tools*, and the resulting latent representation used for clustering and visualisation. Cluster marker genes were identified using the differential gene expression function of *scVI tools* followed by manual annotation based on canonical markers and reference to the literature. Pathway analysis comparing conditions was performed by gene set enrichment analysis with *fgsea* (in R), following pseudobulking of gene expression of the examined cluster with *decoupler* (1.6).

### Statistical analysis

The use of the statistical tests is indicated in the respective figure legends, with the error bars indicating the mean ± S.E.M. P values ≤ 0.05 were considered to indicate significance. Analyses were performed with GraphPad Prism v9-10 or R 4.1. No statistical methods were used to pre-determine sample sizes, but our sample sizes are similar to those reported in previous publications. The distribution of data was determined using normality testing to determine appropriate statistical methodology, or otherwise assumed to be normally distributed. For in vivo experiments we matched the age of the mice in experimental batches, but other modes of randomization were not performed. Data collection and analysis were not performed blind to the conditions of the experiments in most of the experiments. In selected experiments, batch correction was performed with the R package Batchma.

## Acknowledgements

We thank Johanna ten Hoeve-Scott (UCLA Metabolomics Center, United States) for performing cellular metabolomics studies. We thank Tal Arnon for providing X31 influenza (H3N2) strain. We thank Lion Uhl, João Pedro Pereira Bonifacio Lopes, Claire McIntyre, Joy Edwards-Hicks, and Shih-Hsuan Mao for technical assistance and scientific advice. We also thank Jonathan Webber for flow sorting and the BSU staff for support (Kennedy Institute of Rheumatology, University of Oxford). We would like to acknowledge the Oxford-ZEISS Centre of Excellence in Biomedical Imaging and thank its staff. We acknowledge the contribution to this study made by the Oxford Centre for Histopathology Research and the Oxford Radcliffe Biobank, which are funded by the University of Oxford, the Oxford CRUK Cancer Centre, the NIHR Oxford Biomedical Research Centre (BRC) (Molecular Diagnostics Theme/Multimodal Pathology Subtheme and the NIHR CRN Thames Valley network). Oxford Radcliffe Biobank (ORB) research tissue bank ethics, reference 19/SC/0173 for tonsil sections.

## Author contributions

Y.F.Y., E.M., and A.C. conceived and designed the study. Y.F.Y., E.M., K.B., H.A., Z.Y.W, R.M, J.F., J.C.J, M.A, M.B., I.F. and B.K. performed experiments. A.C., Y.F.Y., and E.M. analysed cellular metabolomic and transcriptional data. I.F., J.I.M. and M.D. performed and analysed additional metabolomic experiments. Y.F.Y. and R.M. performed image analysis. C.B., M.C., and S.J.D. provided resources. A.C. and I.R. analysed single cell RNA sequencing data. Y.F.Y. and A.C. wrote the manuscript and A.C. supervised the study.

## Data and material availability

The scRNA sequencing data generated in this study has been deposited to GEO under accession code GSE277338.

